# The spatial arrangement of laminar thickness profiles in the human cortex scaffolds processing hierarchy

**DOI:** 10.1101/2023.03.25.532115

**Authors:** Amin Saberi, Casey Paquola, Konrad Wagstyl, Meike D. Hettwer, Boris C. Bernhardt, Simon B. Eickhoff, Sofie L. Valk

## Abstract

The human neocortex consists of tangentially organized layers with unique cytoarchitectural properties. These layers show spatial variations in thickness and cytoarchitecture across the neocortex, which is thought to support brain function through enabling targeted corticocortical connections. Here, leveraging maps of the six cortical layers in 3D human brain histology, we aimed to quantitatively characterize the systematic covariation of laminar structure in the cortex and its functional consequences. After correcting for the effect of cortical curvature, we identified a spatial pattern of changes in laminar thickness covariance from lateral frontal to posterior occipital regions, which differentiated the dominance of infra- versus supragranular layer thickness. Corresponding to the laminar regularities of cortical connections along cortical hierarchy, the infragranular-dominant pattern of laminar thickness was associated with higher hierarchical positions of regions, mapped based on resting-state effective connectivity in humans and tract-tracing of structural connections in macaques. Moreover, we show that regions with comparable laminar thickness patterns correspond to inter-regional structural covariance, maturational coupling, and transcriptomic patterning, indicating developmental relevance. In sum, here we characterize the association between organization of laminar thickness and processing hierarchy, anchored in ontogeny. As such, we illustrate how laminar organization may provide a foundational principle ultimately supporting human cognitive functioning.

## Introduction

Cortical cytoarchitecture, that is, the organization and characteristics of neurons across the depth of the cortex, varies markedly across the cortical mantle [1–4]. Characterizing this variation has been an important focus of histological studies over the past century. Early studies were largely based on visual inspection and qualitative descriptions of cytoarchitectural features across the cortex to identify local borders between regions [2] or describe more global cytoarchitectural variations [4,5]. With methodological advances of recent decades, there has been a shift towards more quantitative investigations of cortical cytoarchitecture, based on statistical analysis on 2D histological sections [6–8]. The central idea of these studies has been to quantify the variation of cell-body-stained image intensity across the cortical depth, i.e., ‘cortical profile’, followed by observer-independent analysis of how cortical profiles vary across the cortex and define borders of regions, particularly with respect to the central moments, i.e., mean, standard deviation, kurtosis and skewness [1,6,7]. The release of BigBrain, a whole-brain ultra-high-resolution post-mortem histological atlas of a 65 year old male [9], enables such quantitative investigations at a much larger scale, for example to quantify large-scale microstructural gradients at a neocortical [10] and mesiotemporal level [11].

Quantitative studies of cortical profiles have helped improve our understanding of cytoarchitectural variability of the human cortex. However, the cerebral cortex is a layered structure, and models of cortical profiles are, at least explicitly, agnostic to cortical layering. The layers in the neocortex are generally described as six horizontally superimposed stripes of gray matter with characteristic features such as size, type and density of the neurons, which can again be differentiated into multiple sublayers [1,4]. From the pial to the gray-white matter interface, they include layer I which contains mostly dendrites and axon terminals and has a low cellular density, layer II which is densely packed with small granule cells, layer III which contains sparse but large pyramidal neurons, layer IV which consists of densely packed and extremely small granule cells, layer V which is composed of fine pyramidal neurons, and layer VI which includes dense spindle-shaped neurons [2,4]. One of the prominent cytoarchitectural features that vary across the cortex is its laminar structure, with respect to laminar thickness, as well as neuronal size and density of each layer. Indeed, laminar features have been an important focus of many qualitative studies of human cytoarchitectural variation [3,4,12]. For example, agranular and dysgranular cortical types are defined based on the absence or thinness of layer IV, relative to eulaminate and koniocortical regions [4,12]. However, studies on quantitative analysis of cortical cytoarchitecture with respect to its laminar features in humans are limited. Yet, understanding layered organization of the human neocortex may provide further insights into how intracortical circuits ultimately support function [13,14].

According to the ‘structural model’, variation in cortical cytoarchitecture is suggested to scaffold the laminar pattern and strength of inter-regional connections, [3,15]. Specifically, the difference of cytoarchitecture between regions can predict the ‘feedback’, ‘feedforward’ or ‘lateral’ pattern of their laminar connections, and in turn, their relationship along the cortical hierarchy [3,16,17]. For example, recent work using layer-based functional magnetic resonance imaging (fMRI) could show that specific cortical layers are involved in different aspects of memory processing in the dorsolateral prefrontal cortex [18]. Such differences in cognitive processing may be rooted in the connectivity profiles associated with different layer depths. Indeed, the ‘structural model’ proposes that the similarity of cytoarchitecture between two regions can predict a higher likelihood and strength of their connectivity [3,16,19,20]. Here, we aimed to study the organization of laminal profiles across the cortical mantle and its relevance to cortical hierarchy and inter-regional connectivity in the context of the ‘structural model’ to further understand the relationship between human cortical structure and function. To do so we leveraged previously reported maps of the locations of cortical layers across isocortical regions of the BigBrain that were predicted using a convolutional neural network [21]. We extend previous work investigating the spatial arrangement of cortical profiles based on microstructure [10], through formally probing layer-profiles in this model, enabling us to evaluate how the differential arrangement of cortical laminae may contribute to cortical hierarchy and connectivity.

The regional variability of laminar structure and its association to connectivity is suggested to have developmental origins [20,22–24]. For example, the regional variability of laminar neuronal density in the cerebral cortex is suggested to occur due to the regional and laminar differences in neurogenesis timing during early fetal development [20,23,25]. Specifically, while the onset of early fetal neurogenesis is relatively uniform across the cortex, it ends later in regions such as V1 compared to limbic areas. Given upper layers develop later than deeper layers [25], this may explain why regions such as V1 have a higher neuronal density in upper layers [20,23]. The overlap in developmental timing of regions with similar laminar structure can also explain their higher likelihood of connectivity, as neurons born around the same time are shown to have a higher chance of connecting to each other [20,26,27]. Indeed, the shared ontogeny of different cortical regions has been suggested to underlie the systematic covariation of morphometric features in the human cortex, measured across the population. In previous work we could show these patterns vary along posterior-anterior and superior-inferior axes, reflecting functional and evolutionary patterning [28]. However, to what extent these macroscale inter-individual patterns of structural covariation may reflect layer-specific structural arrangement is not yet understood. Therefore, our third aim was to investigate the developmental hypothesis of laminar structure variation.

In this article, we describe a data-driven axis of laminar thickness covariance by quantifying the inter-regional covariation of laminar thickness in the BigBrain [9,21] and evaluate its relation to connectivity as hypothesized by the structural model [3,15]. To do so, we employ dimensionality reduction techniques to identify the principal axis across which laminar thickness covaries. We next evaluate how laminar thickness covariation relates to asymmetry- and laminar-based cortical hierarchy, as well as likelihood and strength of inter-regional connectivity. Last, we focus on the developmental hypothesis of laminar structure variation by evaluating its links to inter-regional structural covariance and maturational coupling, as well as the developmental trajectory of gene expression associated with the laminar structure variation.

## Results

### Principal axis of laminar thickness covariance

We used the maps of cortical layers based on the BigBrain, an ultra-high-resolution post-mortem histological atlas of a 65 year old male [9,21], to study the laminar thickness covariation across the cerebral cortex (Fig. 1a). We first excluded agranular and dysgranular regions, such as cingulate, anterior insula, temporal pole and parahippocampal cortices, in addition to allocortex, given their lack of a six-layer structure [12]. Cortical folding impacts the laminar structure, such that layers inside of the fold are compressed and thicker, whereas layers outside of the fold are stretched and thinner, and this is known as the Bok principle [29–32]. Accordingly, in the BigBrain we observed that from the sulci to the gyri, the relative thickness of superficial layers decreases (r = -0.27, p_spin_ < 0.001) (Fig. S1a). To reduce the local effects of curvature on laminar thickness, we smoothed laminar thickness maps using a moving disk, which reduced this effect remarkably, as the correlation of curvature with the relative thickness of superficial layers dropped to r = -0.12 (p_spin_ < 0.001) (Fig. S1a). Following, the laminar thickness maps were normalized by the total cortical thickness at each cortical location to get the relative thickness. The maps of relative laminar thickness were then parcellated using the Schaefer- 1000 parcellation (Fig. 1b,c). We next calculated the laminar thickness covariance (LTC) matrix, showing the similarity of laminar thickness patterns between cortical areas. The LTC matrix was created by calculating the pairwise partial correlation of relative laminar thickness between cortical locations (controlled for the average laminar thickness across the isocortex), which was subsequently z-transformed (Fig. 1d, Fig. S2).

**Fig. 1.**
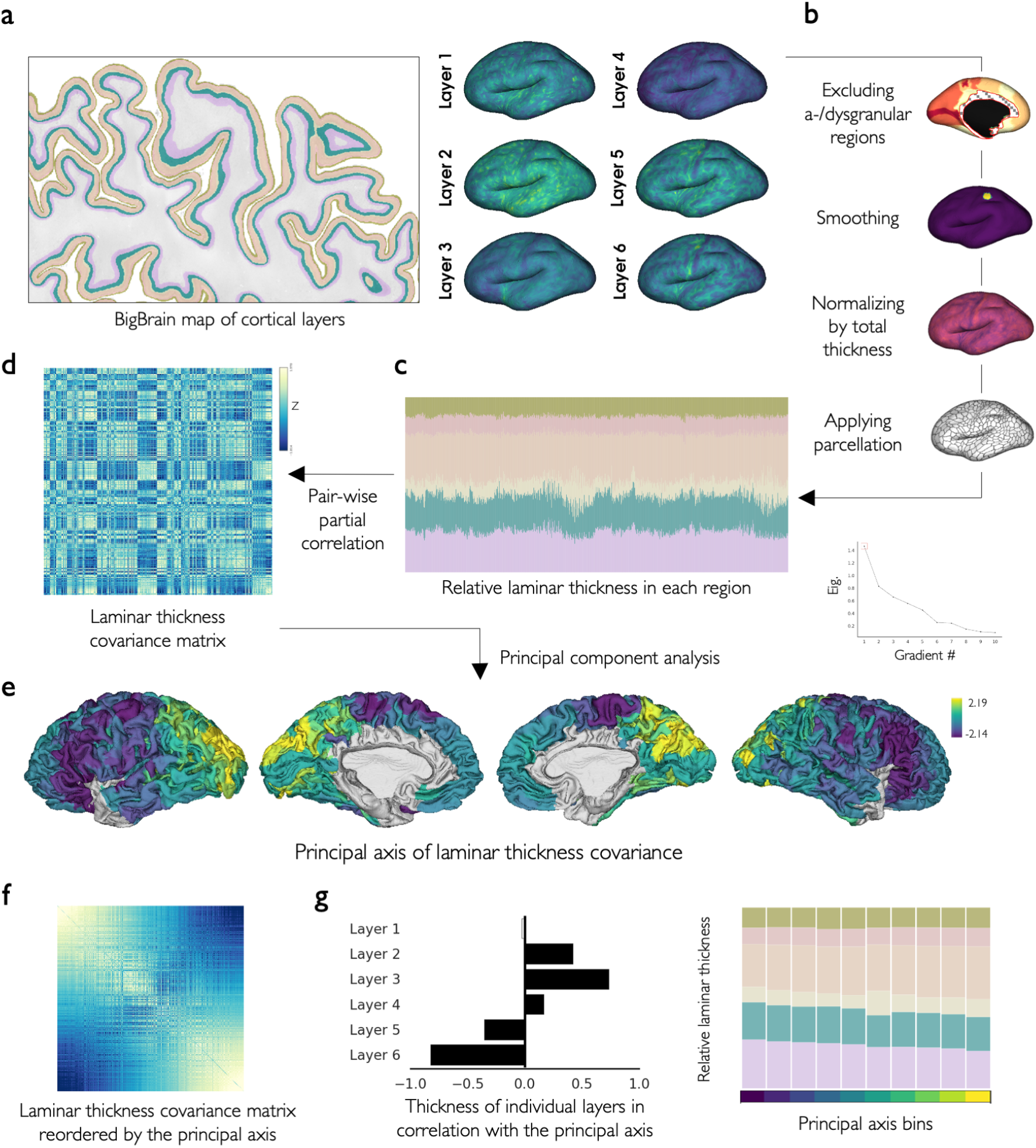
Laminar thickness covariance and its principal axis. **a)** The laminar thickness maps based on the post-mortem histological atlas of BigBrain. **b**,**c)** For each cortical layer, the a-/dysgranular regions were excluded, the thickness map was smoothed using a disc, normalized by the total thickness, and parcellated. **d)** The LTC matrix was created by calculating the pairwise partial correlation of relative thickness across layers and between regions. **e)** The main axis of laminar thickness covariance (LTC G1) was calculated by principal component analysis. **f)** LTC G1 reorders the LTC such that closer regions on this axis have similar LTC patterns. **g)** LTC G1 characterized a shift of infra- to supragranular dominance.

Principal component analysis was then applied to the LTC matrix to identify the axes or gradients along which differences in the loadings indicates regional dissimilarity in the laminar thickness pattern [33]. Here, we focused on the principal axis, LTC G1, which explained approximately 28.1% of the variance in LTC (see the second and third axes in Fig. S3). LTC G1 spanned from the lateral frontal regions, towards medial frontal, temporal and primary visual areas, ending in the parietal and occipital regions (Fig. 1e). This axis was correlated with the relative thickness of layers II (r = 0.42, p_variogram_ < 0.001), III (r = 0.73, p_variogram_ < 0.001) and IV (r = 0.16, p_variogram_ < 0.001) positively, and layers V (r = -0.35, p_variogram_ < 0.001) and VI (r = -0.82, p_variogram_ < 0.001) negatively, characterizing a shift from the dominance of infra- to supragranular layers (Fig. 1g). The spatial map of LTC G1 was mostly robust to analytical choices, i.e., using unparcelled data (20484 vertices) as well as alternative parcellation schemes, covariance metrics, dimensionality reduction techniques, sparsity ratios and the inclusion of a-/dysgranular regions (Fig. S4). In addition, while LTC was higher between physically proximal regions in an exponential regression model (R2 = 0.16, p_spin_ < 0.001), the spatial map of LTC G1 was robust to the effects of geodesic distance (r = 0.97, p_variogram_ < 0.001) (Fig. S5). We also showed that three- and four-layer models of LTC G1 created by merging selected layers were similar to the original six-layer model (Fig. S6). Moreover, as an alternative data-driven approach of quantifying organization of laminar thickness variability K-means clustering was used which revealed four optimal clusters of the regional laminar thickness profiles that were largely aligned with the LTC G1 (F = 813.1, p_spin_ < 0.001) (Fig. S7).

Microstructural profile covariance (MPC) is based on the image intensity profiles in the BigBrain cortex and is a data-driven model of cytoarchitecture that is explicitly agnostic to layer boundaries [10]. MPC was significantly correlated with our model of laminar thickness covariation, at the level of matrices (r = 0.34, p_spin_ < 0.001) and their principal axes (r = 0.55, p_variogram_ < 0.001) (Fig. S8). Further, a model of laminar structure covariation, which took both laminar thickness and laminar density into account, revealed a similar axis significantly correlated with LTC G1 (r = 0.84, p_variogram_ < 0.001) (Fig. S9). In addition, we used a preliminary dataset of layer-wise neuron segmentations, based on higher-resolution 2D patches from select cortical regions of the BigBrain (Fig. S10a), and observed variation of laminar neuronal features along LTC G1 which was most prominent in layer IV, showing nominal increase of neuronal density (r = 0.54) and decrease of neuronal size (r = -0.55) (Fig. S10c,d). In addition, the ratio of average neuronal size in layer III to layer V (externopyramidization) increased along LTC G1 (r = 0.27, Fig. S10e). Last, comparing our data-driven model of laminar thickness covariation with the map of cortical types, a theory-driven model of laminar structural variation [12], we observed no significant association (F = 6.41, p_spin_ = 0.633) (Fig. S11).

### Principal axis of laminar thickness covariance in association with cortical hierarchy

After describing how the overall laminar thickness pattern varies across the cortex, we sought to understand its relation to the asymmetry- and laminar-based cortical hierarchy.

Asymmetry-based hierarchy was defined based on the group-averaged whole-brain effective (directed) connectivity of brain regions. The effective connectivity matrix (Fig. 2a) shows the influence of each brain region on the activity of other regions, and was previously estimated using regression dynamical causal modelling (rDCM), based on resting-state fMRI data from 40 healthy adults [34–37]. Using the effective connectivity matrix, we calculated the asymmetry-based hierarchy of each region as the difference between its weighted out-degree (efferent strength) and in-degree (afferent strength), showing to what extent it influences the activity of other regions as opposed to being influenced by them. The asymmetry-based hierarchy map was significantly correlated with LTC G1 (r = -0.40, p_variogram_ < 0.001), indicating higher hierarchical position of infragranular-dominant regions (Fig. 2b). Accordingly, the asymmetry-based hierarchy map was significantly correlated with the relative thickness of layers III and IV negatively, and layers V and VI positively (Fig. S12). Of note, decomposing the asymmetry-based hierarchy into its components, we observed a significant correlation of LTC G1 with the weighted in-degree (r = 0.61, p_variogram_ < 0.001) but not out-degree (r = -0.01, p_variogram_ = 0.868) (Fig. S13). The asymmetry-based hierarchy map calculated using a replication sample based on the Human Connectome Project (HCP) dataset (N = 100) [36,38] was similarly correlated with LTC G1 (r = -0.49, p_variogram_ < 0.001; Fig. S14). Note that for the above analyses, we recalculated LTC and LTC G1 in the Schaefer-400 parcellation, as the effective connectivity matrices obtained from the previous work by Paquola and colleagues [36] was available in this parcellation.

**Fig. 2.**
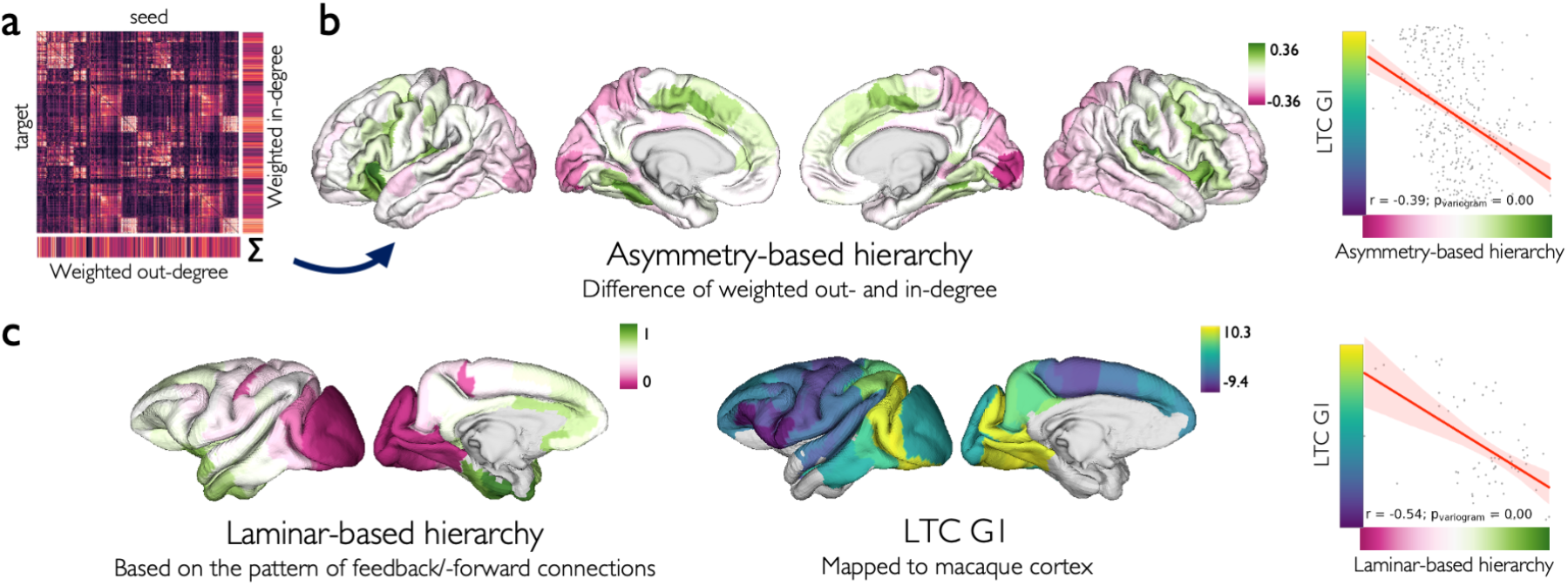
Association of laminar thickness covariance and cortical hierarchy. **a)** The group-averaged effective connectivity matrix based on regression dynamic causal modeling. **b)** Regional asymmetry-based hierarchy was calculated as the difference between their weighted unsigned out-degree and in-degree, and was significantly correlated with the LTC G1. **c)** Regional laminar-based hierarchy map of macaque was correlated with the LTC G1 aligned to the macaque cortex.

In addition, we obtained the laminar-based hierarchy map of the macaque cortex from a previous study [39]. Laminar-based hierarchy assumes higher hierarchical positions for regions projecting feedback and receiving feedforward connections, as quantified in tract-tracing studies. This approach may be a more direct measure of processing hierarchy in terms of laminar regularities of connections [39] but is not available in humans. After aligning the LTC G1 map of the human cortex to the macaque’s cortex [40], we observed that it was significantly correlated with the map of macaque’s laminar-based hierarchy (r = -0.54, p_variogram_ < 0.001; Fig. 2c). In addition, laminar-based hierarchy showed significant positive correlations with the relative thickness of layers III and IV, and negative correlations with the relative thickness of layers V and VI (Fig. S12). These findings indicated association of laminar thickness variation to two alternative maps of cortical hierarchy based on asymmetry of effective connectivity and laminar pattern of cortical connections.

The regional distribution of excitatory and inhibitory neuronal subtypes have been suggested to vary with cortical hierarchy and laminar structure [23,39,41]. Thus, we studied this relationship with our laminar covariance axis using gene expression data obtained from the Allen Human Brain Atlas (AHBA; N = 6) [42,43] and the list of genes associated with each neuronal subtype (Table S1) [44]. We observed significant FDR-corrected correlations of LTC G1 with 11, asymmetry-based hierarchy with 10 and laminar-based hierarchy with 14 out of 16 neuronal subtypes (Fig. S15).

### Laminar thickness covariance in association with cortico-cortical connectivity

After showing alignment of asymmetry- and laminar-based hierarchy with laminar thickness variation, we studied whether the similarity of regions in laminar thickness relates to the likelihood and strength of interareal connectivity (Fig. 3).

**Fig. 3.**
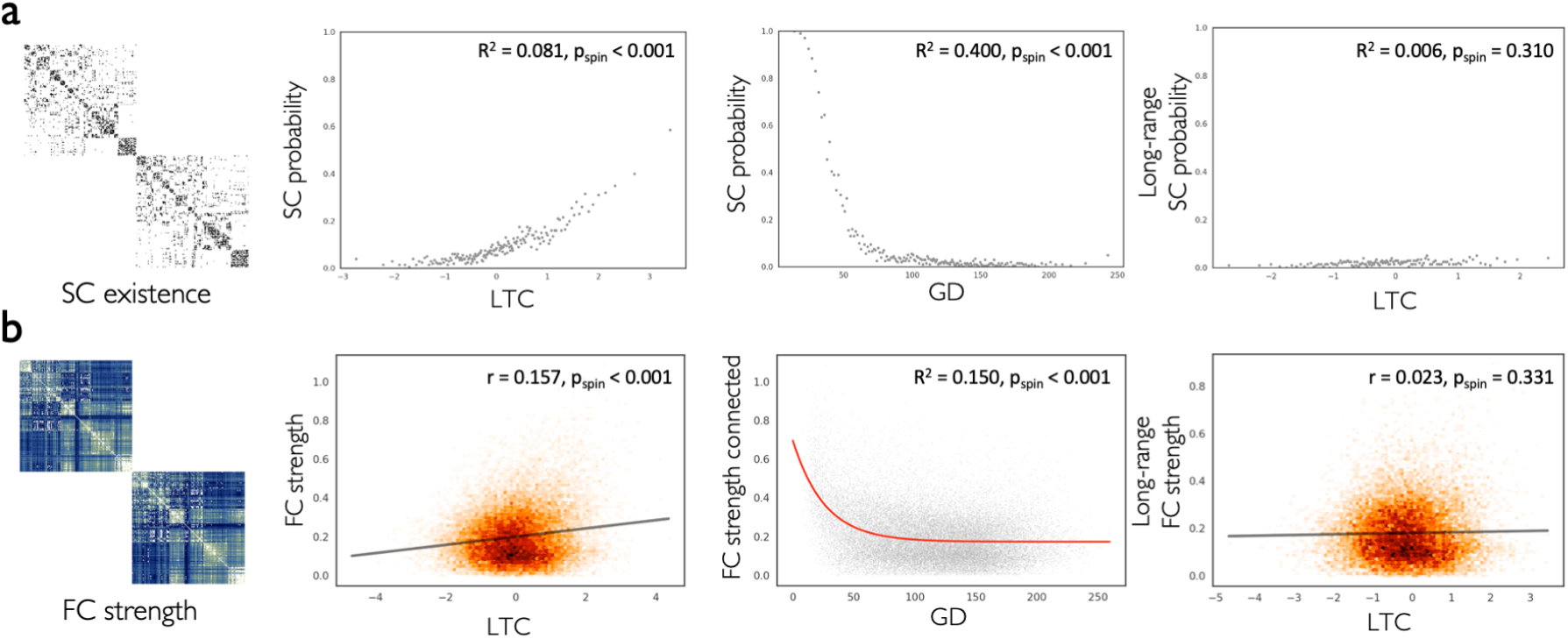
Association of laminar thickness covariance with connectivity. **a)** The binarized structural connectivity (SC) matrix showing the existence of intra-hemispheric connections (left). SC likelihood was associated with increased laminar thickness covariance (LTC) (center left) and decreased geodesic distance (GD) (center right). SC likelihood among long-range connections was not significantly associated with LTC. **b)** The functional connectivity (FC) matrix showing the strength of intra-hemispheric connections (left). FC strength was correlated with increased LTC (center left) and decreased exponentially with GD (center right). FC strength among long-range connections was not significantly correlated with LTC.

We used the structural and functional connectivity (SC and FC) matrices (400 regions) averaged across a subgroup of the HCP dataset (N = 207) [38,45], which was obtained from the ENIGMA (Enhancing NeuroImaging Genetics through Meta-Analysis) Toolbox [46]. Using logistic regression, we observed higher LTC was associated with the increased likelihood of SC (R2 = 0.082, p_spin_ < 0.00). In addition, LTC was correlated with the increased strength of FC (r = 0.15, p_spin_ < 0.001). Previous research indicates that neighboring regions in the cerebral cortex are more likely to connect [47,48] and also tend to have similar structural and functional features [49,50]. Here we also observed this effect, with physically proximal regions showing higher likelihood of SC (R2 = 0.400, p_spin_ < 0.001) and strength of FC (R2 = 0.150, p_spin_ < 0.001) on one hand, and higher LTC (R2 = 0.164, p_spin_ < 0.001) on the other hand. To understand whether LTC was associated with connectivity independent of these distance effects, we studied the association of LTC with long-range connectivity. We observed that LTC was not significantly associated with the likelihood of long-range SC (R2 = 0.006, p_spin_ = 0.310), or strength of long-range FC (r = 0.023, p_spin_ = 0.331). This finding suggested inter-regional distance as an important covariate in the association of LTC with connectivity.

### Maturational and transcriptomic links to laminar thickness covariance

Thus far, we described how the overall laminar structure varies across the cortex and its relevance to cortical hierarchy and connectivity. Lastly, we sought to study the question of potential links of laminar thickness variation to maturational and genetic factors.

Structural covariance, i.e., the pattern of covariation in cortical morphology (e.g., cortical thickness) across a population, provides a model of shared developmental/maturational and genetic effects between cortical regions [28,51,52]. We obtained the structural covariance matrix based on the HCP S1200 release (N = 1206) from our previous work [28], and observed that it was significantly correlated with the LTC at the level of matrices (r = 0.33, p_spin_ < 0.001) and their principal axes (r = -0.57, p_variogram_ < 0.001). This suggests shared developmental/maturational and genetic effects between regions with similar laminar thickness (Fig. 4a). Decomposing structural covariance into genetic (heritable) and environmental components [28], we observed correlation of LTC with the inter-regional genetic correlation (r = 0.30, p_spin_ < 0.001) (Fig. S16). Furthermore, focusing on the developmental effects, we obtained the interregional maturational coupling matrix from a previous study by Khundrakpam and colleagues [52]. This matrix shows the similarity of regions in longitudinal cortical thickness changes over development in a dataset of children and adolescents (N = 140, baseline age = 11.9±3.6, followed up for 2 years), and was weakly correlated with the LTC matrix (r = 0.10, p_spin_ < 0.001) (Fig. 4b).

**Fig. 4.**
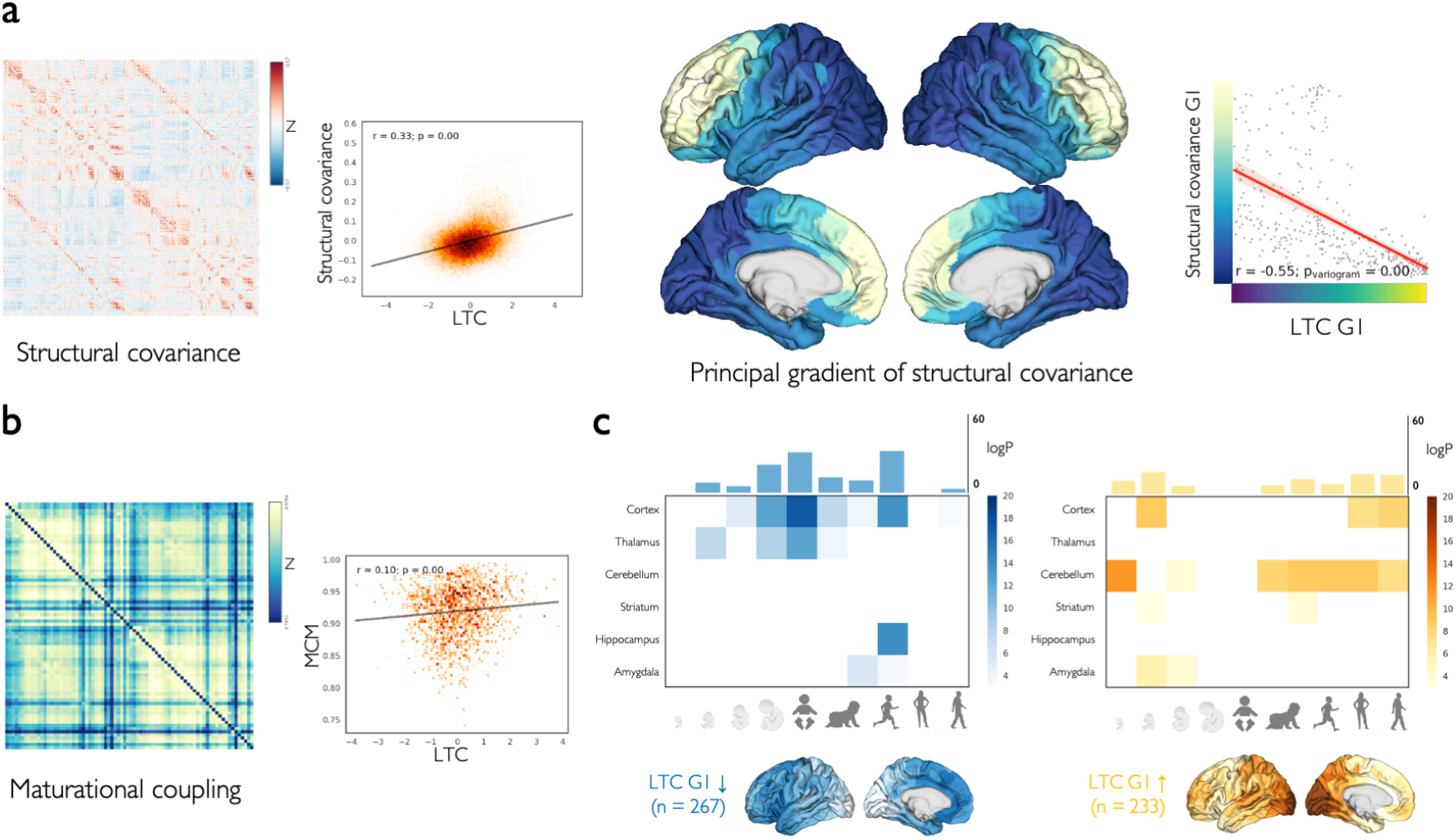
Maturational and transcriptomic links to laminar thickness covariance. **a)** The structural covariance matrix based on cortical thickness (left) in association with the LTC (center left). Main axes of structural covariance (center right) and LTC were correlated (right). **b)** Maturational coupling matrix (left) was weakly associated with LTC (right). **c)** The average expression of top genes with regional expressions aligned to LTC G1, over-expressed rostrally (blue) or caudally (yellow), and their spatiotemporal developmental enrichment.

A second approach to investigate potential covariation of laminar structure as a function of development and genetics was done by using brain transcriptomics. Further supporting the idea of shared genetic effects between regions with similar laminar thickness, we first showed a correlation of r = 0.20 (p_spin_ = 0.010) between LTC and the correlated gene expression matrix. This matrix shows the correlation of gene expression between regions and across all genes, based on transcriptomics data obtained from the AHBA (Fig. S17) [42,43]. Next, we identified distinct sets of genes expressed preferentially at the opposite ends of the LTC G1 in adulthood based on the AHBA data, and investigated their developmental enrichment using the BrainSpan dataset [53]. We observed distinct spatiotemporal patterns for the expression of each set of genes associated with the rostral versus caudal ends of the LTC G1. Specifically, within the cerebral cortex, genes preferentially expressed at the caudal end of LTC G1 are mainly expressed in mid fetal and post-pubertal stages, while genes with higher expression at the rostral end of LTC G1 were expressed in late fetal and pre-pubertal stages of development (Fig. 4c).

## Discussion

In the current study, we sought to extend previous quantitative studies on cytoarchitectural variability of the cerebral cortex [1,10], by focusing on the layered structure of the cerebral cortex. We used the map of cortical layers [21] based on the ultra-high-resolution atlas of BigBrain [9] to identify a principal axis of laminar thickness covariation in the cortex. We observed an axis of laminar thickness covariance showing a shift from the dominance of supragranular towards infragranular layers thickness from occipital to lateral frontal areas. This shift was co-aligned with cortical hierarchy, defined either based on the asymmetry of afferent and efferent connections in the human cortex, or the laminar pattern of connections in the macaque cortex. We also found a higher likelihood and strength of connections between regions with similar patterns of laminar thickness, supporting the principle of “similar prefers similar” in cortical wiring and the structural model of connectivity [3,15,16,20]. Finally, we showed that laminar thickness covariation is linked to the inter-regional morphological covariance and transcriptomic expression variability across development, supporting the prevailing ideas on the potential origins of laminar structure variability [20,22,23].

The principal axis of laminar thickness covariation characterized an overall increase in the relative thickness of supragranular layers from the lateral frontal to posterior occipital regions. This was in line with a previous animal study which illustrated relative increase in the implied column height of the upper layers along the rostro-caudal axis of the cortex in several rodent and non-human primate species [22]. The same study also reported that from rostral to caudal regions the density of neurons increases, as had been shown in a few other studies [54–56], but the increase selectively occurs in the layers II-IV. Using a preliminary dataset of laminar neuronal features in a few cortical regions of the BigBrain, we also observed an increased neuronal density along LTC G1 which was most prominent in layer IV. In addition, we observed increased size of supra- versus infragranular neurons, which may indicate externopyramidization along this axis. Of note, these analyses should be interpreted with caution as they were limited to a small number of available samples, making proper statistical tests not applicable. Overall, from the lateral frontal towards parietal and occipital regions, the granular and supragranular cortical layers become more prominent relative to the infragranular layers, with respect to thickness, and potentially neuronal density and soma size.

Previous theory-based approaches based on visual inspection of histological samples have additionally described a sensory-fugal axis of laminar structure variation transitioning from sensory to limbic regions [4,12]. This qualitatively observed sensory-fugal axis is overall different from the quantitative axis of laminar thickness covariation that we described, which may be attributed to the different features of interest studied, such as the prominence of layer IV, as opposed to laminar thickness (and gray matter intensity) in our study. We observed that regions that are part of the same cortical type show variations with respect to the laminar thickness pattern, which may indicate differential processes underlying laminar structure. In fact, a previous data-driven model of microstructural profile covariance in the BigBrain revealed two main axes of cytoarchitectural variability: a rostro-caudal and a sensory-fugal axis [10]. These different axes may reflect diverging neurobiological routes organizing the human cortex. Indeed, in our previous work on group-level cortical thickness covariance and genetic correlations, we observed a a rostral-caudal axis which was suggested to reflect differentiation between cortical hierarchy and maturational effect, and a ventral-dorsal axis reflecting microstructural pattern associated with the theory of dual origin [28,57]. Thus overall it is likely that multiple neurobiological axes organize the human cortex, reflecting distinct maturational and (phylo)genetic effects.

In the current work, we focused on layer-thickness averaged in 400-1000 functional parcels. There has been some debate over the optimal approach and level of granularity to study cytoarchitectural variability of the cortex [1,58]. Previous studies have ranged from focusing on fine cytoarchitectural details and identification of sharp borders between regions [2,4] to classification of the cortex into broader categories with grossly comparable cytoarchitecture [4,12]. On the other hand, some authors have argued against cortex-wide existence of sharp boundaries and rather focused on the more gradual variations across the cortex [5,58]. We should note that here we refrained from making any assumptions on the (non)existence of sharp borders or a level of granularity as we aimed to provide a whole-cortex layer covariance organizational axis. We argue that the topology of cytoarchitectural variability of the cortex ranges from abrupt to more gradual changes [1,58]. Accordingly, the LTC G1 map consisted of a combination of sharp borders and gradual transitions, but was more dominated by gradual changes. However, additional sharp borders emerge in the subsequent principal components, and indeed, we should not expect to identify all regional borders solely based on a single measure, i.e., relative laminar thickness [1]. Future work at 1 micrometer resolution images of the BigBrain may further uncover the organization of borders and abrupt alterations of layer thickness and associated cytoarchitecture variation.

The main axis of laminar thickness covariance aligned with the asymmetry- and laminar-based cortical hierarchy, extending previous observations on the link between cortical hierarchy and microstructure [17,39]. Specifically, regions towards the infragranular-dominant end of the axis were positioned higher in the cortical hierarchy than the supragranular-dominant regions. The link between laminar thickness and cortical hierarchy can be explained in the context of regularities in the laminar pattern of feedforward and feedback inter-areal connections along the cortical hierarchy, as reported in tract-tracing studies on the mammalian cerebral cortex [59–61]. Feedforward connections transmit high-dimensional information up the processing hierarchy and originate from the supragranular layers II and III, whereas feedback connections propagate abstract representations down the hierarchy and originate from infragranular layers V and VI [59–63]. Therefore, along the cortical hierarchy, there is an increase in the dominance of feedback processing, and as the infragranular layers are the source of feedback connections, it is not surprising to find increased infragranular thickness in regions further up in the hierarchy. Of note, several definitions exist for cortical hierarchy, and alternative mappings based on theory [63] or proxy measures, such as the principal axis of functional connectivity [64], or T1w/T2w ratio [39] have been used in the literature. The mapping of cortical hierarchy may vary based on its definition [65] or the experimental condition, e.g., resting state in contrast to various tasks. We defined cortical hierarchy based on asymmetry of resting-state effective connectivity in humans as well as laminar pattern of connections in macaques. The asymmetry-based hierarchy assumes that regions higher up in the processing hierarchy have stronger efferent than afferent connections, that is, they influence the activity of lower regions rather than being influenced by them [65–67]. The hierarchy in macaque was defined based on the laminar pattern of feedforward and feedback connections identified in tract-tracing studies [39], providing a more direct measurement.

Furthermore, we observed that LTC G1 and the maps of cortical hierarchy were co-aligned with regional variability of several subtypes of excitatory and inhibitory neurons, mapped based on aggregated expression of their associated genes [39,42,44]. These neuronal subtypes are localized across different cortical layers [44,68] and were previously shown to correlate with an alternative mapping of cortical hierarchy based on T1w/T2w ratio [39]. In addition, varying density of parvalbumin-positive and calbindin-positive inhibitory neuronal subtypes along cortical types is reported in the macaque cortex [23]. Overall, these findings suggest a gradient of cortical microcircuitry which is aligned with cytoarchitectural variability of the cortex and cortical hierarchy [41]. This gradient of microcircuitry has important functional implications [41]. For example, along the cortical hierarchy from V1 to the prefrontal cortex there is an increase in the density of dendritic spines, which is suggested to lead to higher strength of recurrent connections, which in turn, is necessary for persistent neural activity in the prefrontal needed for working memory [41]. Furthermore, incorporating regional heterogeneity of excitation and inhibition [69] or microstructure (T1w/T2w ratio) [69–71] in the dynamical models of the brain improves the model fit to the empirical resting-state fMRI data.

The similarity of regions in their laminar thickness patterns was associated with an increased likelihood and strength of their connectivity. This finding supported the structural model for connectivity relating cytoarchitectural similarity to the likelihood and strength of connections [3,16,19], and was in line with studies showing higher likelihood and strength of connections between regions with similar microstructure, based on the complexity of pyramidal neurons [72], neuronal density [20,73] or cortical types [19,20,26,74]. In addition, and of particular relevance to our findings, interareal connectivity in the human cortex has been linked to the microstructural profile covariance of the BigBrain [75–77]. Specifically, connected regions were reported to have higher similarity in their microstructural profiles compared to non-connected profiles, and microstructural profile covariance correlated with the connectivity strength [77]. Furthermore, a previous study used generative modeling of connectivity and showed that including both microstructural profiles covariance and wiring cost in the model, as opposed to including wiring cost alone, leads to a better fit [75]. In addition, a low-dimensional coordinate space of the human cortex calculated by incorporating inter-regional structural connectivity, physical proximity and the BigBrain’s microstructural covariance was shown to predict functional connectivity with a high accuracy [76]. In our study we extend these findings and show that the probability and strength of connectivity additionally relates to the laminar thickness profiles in the BigBrain. Using an alternative approach, another recent study focused on the interrelation between connectivity and the absolute thickness of individual layers in the BigBrain, and showed that regions with thicker layer IV are less likely to connect to regions with higher thickness in layers III, V and VI [78]. Overall, these findings are in line with the wiring principle of “similar prefers similar” [20,26,75,78], which has been observed not only with the similarity of microstructure, but also in association to gene expression patterns [79–83], neurotransmitter receptor profiles [1] and macroscale morphometry [78,85]. It may be argued that this ubiquitous finding simply reflects the fact that nearby cortical regions tend to be similar [49,50], and also are more likely to connect, due to the principle of wiring cost reduction [47,48]. However, this principle alone does not fully account for the connectome architecture, and in particular the existence of long-range connections. For example, simulations were shown to better account for the connectome when inter-regional similarity was considered in addition to the wiring cost reduction [75]. In our study we showed that strength and likelihood of long-range connections were not associated with laminar thickness covariance. A decreased association of microstructural similarity and connectivity among long-range connections has been also observed in previous studies [77]. This raises the question how long-distance connections are encoded in layer-based architecture of the human cortex. Possibly the uncoupling of layer-similarity and long distance connections could be in part driven by an uncoupling through activity-dependent organization, linked to the tethering hypothesis [86]. Further work integrating connectivity with layer-based approaches may help to further understand the interrelationship between short- and long-distance connections and cortical architecture.

The inter-regional similarity of laminar structure and its functional implications have been suggested to arise due to the similar developmental trajectories of the regions [20,22,23]. Here, we observed findings in favor of this hypothesis by showing the correlation of laminar thickness covariance and the population-level inter-regional structural covariance [28,51] as well as subject-level longitudinal maturational coupling of cortical regions [52]. We further used developmental enrichment analysis of the genes overexpressed at the two ends of LTC G1 and showed distinct spatiotemporal trajectories of these genes through pre- and postnatal development, suggestive of developmental timing differences of regions with variable laminar structure. It should be noted however that the transcriptomics data used in the association with LTC G1 was based on post mortem adult brains. Yet, gene expression is dynamic through development [87], and a post mortem snapshot in the adulthood might actually differ from what we would have observed if we had studied the regional variation of gene expression in each developmental stage. Furthermore, the laminar structure changes during development. For example, in the primary motor area, the layer IV which is prenatally prominent, with the maturation of pyramidal neurons in adjacent layers III and V becomes less prominent in the first postnatal months [88]. The underlying mechanisms of how and at which stage during development the adult pattern of laminar thickness covariance may arise remains an open question. Developmental mechanisms such as differences in neurogenesis timing during fetal development [20,22,23], or region- and layer-specific neuronal death in early postnatal stages [24] have been proposed as examples of how regional variations in laminar structure develop. Further studies using postmortem histology or in-vivo markers of laminar structure [10,89,90] are needed to characterize maturation of laminar structure at different stages of development and better understand the underlying mechanisms of regional variations in laminar structure.

### Limitations

There are various limitations that are important to acknowledge in the context of the current work. First, a whole-brain map of cortical layers is currently only available for a single individual, and until a similar atlas becomes available, it is unclear how much the findings presented here would generalize to the other individuals. In addition, the degree to which laminar structure varies across individuals, how it might relate to behavior and function, and its changes through development remain unknown, which highlights the importance of future studies on in-vivo estimation of laminar structure based on high-resolution imaging. Second, while the laminar thickness covariation was evaluated on the BigBrain, it was contextualized in relation to the connectivity, cortical hierarchy and gene expression data that was obtained from different individuals, and this may lead to underestimating the effects due to the potential interindividual variability in these measures. Third, we benefited from using a data-driven approach and a whole-brain map, but in doing so, we only focused on the gross laminar features including thickness and the average cell-body-stained image intensity. On the other hand, theory-driven maps of laminar structure such as cortical types are determined based on a variety of different laminar features [12], yet some of the finer features such as the properties of individual neurons were invisible to our model. Fourth, we studied laminar thickness covariance using a six-layer model of the cortex. However, it is a matter of debate whether all cortical regions have all the six layers, and it has been argued that some regions may have more or fewer number of layers [1,4]. Further work may address this open question with more fine-grained models of intra-cortical structure and metrics to evaluate their functional relevance. This would enable formally testing the optimal architecture of cortical depth. Finally, according to the Bok principle, cortical folding is an important confound in studies of laminar thickness [29–32]. We reduced this effect by smoothing the laminar thickness maps, thereby averaging over local differences of laminar thickness between adjacent sulci and gyri. However, this approach did not completely remove the folding effect, and it may have smoothed over local variations of laminar thickness unrelated to cortical folding, in particular at borders of regions.

### Conclusions

In sum, we described an axis of laminar thickness covariation in the BigBrain, which characterized a structural shift from supra- to infragranular layer thickness. This shift was co-aligned with the cortical processing hierarchy, with infragranular-dominant regions positioned higher across the hierarchy. In addition, regional variation of laminar thickness in the cortex was related to connectivity as well as maturational and genetic patterning of the cortex. Future work may help further understand the relevance of laminar structural variation to human brain function across the lifespan, ultimately providing insights into how the anatomy of the human brain supports human cognition.

## Methods

### BigBrain maps of laminar thickness

BigBrain is a three-dimensional histological atlas of a postmortem human brain (male, 65 year), which is created by digital reconstruction of ultra-high-resolution sections (20 µm) stained for cell bodies, and is publicly available at https://ftp.bigbrainproject.org/ [9]. The cerebral cortex of the BigBrain was previously segmented into six layers, using a convolutional neural network trained on the samples manually segmented by expert anatomists [21]. We used the BigBrain laminar thickness data in the bigbrain surface space, which was included in the BigBrainWarp toolbox (https://bigbrainwarp.readthedocs.io) [91]. The BigBrain surface mesh and laminar thickness maps were downsampled from 164k to 10k points (vertices) per hemisphere to reduce the computational cost of the analyses. The surface mesh was downsampled by selecting a reduced number of vertices and re-triangulating the surface while preserving the cortical morphology, and the surface data (e.g., laminar thickness) were downsampled by assigning the value of each maintained vertex to the average value of that vertex and its nearest removed vertices [91].

### BigBrain layer-specific distribution of neurons

Layer-specific neuronal density and size in selected tissue sections of the BigBrain isocortex at the resolution of 1 µm was obtained from the Python package siibra (https://siibra-python.readthedocs.io/en/latest). This dataset was created by manual annotation of cortical layers and automatic segmentation of neuronal cell bodies (https://github.com/FZJ-INM1-BDA/celldetection) [92]. It contains the data for 111 tissue sections, corresponding to 80 vertices on the BigBrain downsampled surface.

### Laminar thickness covariance

We first excluded regions of the brain with the agranular or dysgranular cortical type due to the less clear definition of the layer boundaries in these regions [12]. The map of cortical types was created by assigning each von Economo region [93] to one of the six cortical types, including agranular, dysgranular, eulaminate I, eulaminate II, eulaminate III, and koniocortex, based on manual annotations published by García-Cabezas and colleagues [12]. Next, for each individual layer and in each hemisphere, the thickness maps were smoothed using a moving disk with a radius of 10 mm to reduce the local effects of curvature on laminar thickness. Specifically, the cortical surface mesh was inflated using FreeSurfer 7.1 (https://surfer.nmr.mgh.harvard.edu/) [94], and for each vertex a disk was created by identifying its neighbor vertices within a Euclidean distance of 10 mm on the inflated surface, and a uniform average of the disk was calculated as the smoothed laminar thickness at that vertex. Next, to obtain the relative laminar structure at each vertex, laminar thicknesses were divided by the total cortical thickness. Finally, the maps of laminar thickness were parcellated using the Schaefer 1000-region atlas [95], of which 889 regions were outside a-/dysgranular cortex and were included in the analyses. The parcellation was performed by taking the median value of the vertices within each parcel. Alternative parcellation schemes including the Schaefer 400-region [95], Desikan-Killiany (68 regions) [96], AAL (78 cortical regions) [97], AAL subdivided into 1012 regions [98] and the Human Connectome Project Multi-Modal Parcellation (180 regions) [99] were used to show robustness of findings, and to enable associations of LTC with the data available in specific parcellations. Of note, the parcellation maps that were originally in the fsaverage space were transformed to the bigbrain space, based on multimodal surface matching and using BigBrainWarp [91,100]. One parcellation (AAL) was originally available in the civet space and was transformed to fsaverage using neuromaps [101] before being transformed to the bigbrain space. In addition to the different parcellation schemes, to further evaluate robustness of our findings to the effect of parcellation the analyses were also repeated on unparcellated data at the level of vertices.

Laminar thickness covariance between the cortical regions was calculated by performing pairwise partial correlation of relative laminar thicknesses, while controlling for the average laminar thickness across the whole cortex, to identify greater-than-average covariance. The partial correlation coefficients were subsequently Z-transformed, resulting in the laminar thickness covariance matrix. We also used alternative covariance metrics including full Pearson’s correlation as well as Euclidean distance in the robustness analyses.

### Geodesic distance

The geodesic distance between two points on the cortical surface refers to the length of the shortest path between them on the mesh-based representation of the cortex. Using the Connectome Workbench (https://www.humanconnectome.org/software/connectome-workbench) [102], we calculated the geodesic distance between the centroids of each pair of parcels, where the centroid was defined as the vertex that has the lowest sum of Euclidean distance from all other vertices within the parcel. The geodesic distance calculation was adapted from its implementation in micapipe (https://micapipe.readthedocs.io) [103]. To evaluate whether our findings were robust to the effect of geodesic distance, in some analyses the effect of geodesic distance on LTC was regressed out using an exponential regression.

### Cortical folding

Mean curvature was calculated at each vertex of the mid-cortical surface based on the Laplace-Beltrami operator using pycortex (https://gallantlab.github.io/pycortex/) [104]. To compute the curvature similarity matrix, for each pair of parcels we estimated their similarity in the distribution of mean curvature across their vertices. This was achieved by calculating Jensen-Shannon divergence of their respective probability density functions.

### Microstructural profile covariance and laminar density covariance

The image intensity of the cell-body-stained BigBrain atlas reflects neuronal density and some size, and its variation across cortical depth at each cortical location is referred to as ‘cortical profile’ or ‘microstructural profile’. We obtained the microstructural profiles sampled at 50 equivolumetric surfaces along the cortical depth from the BigBrainWarp toolbox [91] and reproduced the histological microstructural profile covariance (MPC) matrix as previously done by Paquola and colleagues [10,91]. The microstructural profiles were first parcellated by taking the median. Subsequently, MPC matrix was calculated by performing pairwise partial correlations of regional microstructural profiles, controlled for the average microstructural profile across the whole cortex.

In addition, we created layer-specific cortical profiles by sampling the BigBrain image intensity at 10 equivolumetric surfaces along the depth of each layer, which were then averaged, creating six laminar density maps. Subsequently, a laminar density covariance matrix was created similar to the approach described above for the LTC and MPC matrices.

### Effective connectivity

We obtained the effective connectivity matrix from a prior study [36] based on the microstructure informed connectomics (MICs) cohort (N = 40; 14 females, age = 30.4±6.7) [37] as well as a replication sample from the minimally preprocessed S900 release of the HCP dataset (N = 100; 66 females, age = 28.8±3.8) [38,45]. The effective connectivity matrix between Schaefer 400 parcels was estimated based on rs-fMRI scans using regression dynamic causal modelling [34,35], freely available as part of the TAPAS software package [105]. This approach is a computationally highly efficient method of estimating effective, directed connectivity strengths between brain regions using a generative model.

The effective connectivity matrix was used to estimate asymmetry-based hierarchy, which assumes that hierarchically higher regions tend to drive the activity in other regions rather than their activity being influenced by them. Therefore, given the effective connectivity matrix, after converting it to an unsigned matrix, we calculated the regional asymmetry-based hierarchy as the difference of the weighted out-degree and in-degree of each region, assuming higher hierarchical position for regions with higher efferent than afferent strength.

### Macaque map of cortical hierarchy

The macaque map of laminar-based hierarchy was obtained from a previous work by Burt and colleagues [39]. Briefly, this map was created by applying a generalized linear model to the laminar projection data, based on the publicly available retrograde tract-tracing data (http://core-nets.org) [106], resulting in hierarchy values in 89 cortical regions of macaque’s M132 parcellation [107–109]. To compare the macaque cortical map of laminar-based hierarchy to the human maps of LTC G1 and thickness of individual layers, we aligned these maps to the macaque cortex using the approach developed by Xu and colleagues [40]. Specifically, we first transformed the unparcellated human maps from the bigbrain space to the human fs_LR space using BigBrainWarp [91], mapped it to macaque fs_LR space using the Connectome Workbench, and finally parcellated the transformed map in macaque fs_LR space using M132 parcellation.

### Structural and functional connectivity

The group-averaged functional and structural connectivity matrices based on a selected group of unrelated healthy adults (N = 207; 124 females, age = 28.7±3.7) from the HCP dataset [38,45] were obtained from the publicly available ENIGMA Toolbox (https://github.com/MICA-MNI/ENIGMA) [46]. We fetched the connectivity matrices created in the Schaefer 400 parcellation. We refer the reader to the ENIGMA Toolbox publication and online documentations for the details on image acquistion and processing. Briefly, the functional connectivity matrix for each subject was generated by computing pairwise correlations between the time series of all cortical regions in a resting-state fMRI scan, which after setting negative correlations to zero and Z-transformation were aggregated across the participants. The structural connectivity matrices were generated from preprocessed diffusion MRI data using tractography performed with MRtrix3 [110] and were group-averaged using a distance-dependent thresholding procedure.

### Structural covariance, genetic correlation and environmental correlation

We obtained the structural covariance matrix, as well as the inter-regional genetic and environmental correlation matrices from our previous work [28]. The structural covariance was based on the cortical thickness values of the Schaefer-400 parcels in each individual and was computed by correlating the cortical thickness values between regions and across HCP participants (N = 1206; 656 females, age = 28.8±3.7) [38,45], while controlling for age, sex, and global thickness. Twin-based bivariate polygenic analyses were then performed to decompose the phenotypic correlation between cortical thickness samples to genetic and environmental correlations.

### Maturational coupling

We obtained the group-averaged matrix of maturational coupling from a previous work by Khundrakpam and colleagues based on a sample of children and adolescents (N = 141; 57 females, age at baseline = 11.9±3.6) who were scanned 3 times during a 2-year follow-up [52]. In this study, subject-based maturational coupling was calculated between 78 cortical regions of the AAL parcellation as their similarity in the slope of longitudinal changes in cortical thickness across 3 time points. Subject-based maturational coupling matrices were subsequently pooled into a group-averaged matrix.

### Regional gene expression maps

Regional microarray expression data were obtained from 6 post-mortem brains (1 female, age = 24.0–57.0) provided by the Allen Human Brain Atlas (AHBA; https://human.brain-map.org) [42]. Data was processed with the abagen toolbox (https://github.com/rmarkello/abagen) [43] using the Schaefer-400 surface-based atlas in MNI space.

First, microarray probes were reannotated using data provided by and colleagues [49]; probes not matched to a valid Entrez ID were discarded. Next, probes were filtered based on their expression intensity relative to background noise [111], such that probes with intensity less than the background in ≥ 50% or 25% of samples across donors were discarded. The threshold of ≥ 50% was used for correlated gene expression and developmental gene enrichment analyses, and ≥ 25% was used for creating transcriptomics maps of neuronal subtypes, as it led to removal of fewer genes associated with these subtypes. When multiple probes indexed the expression of the same gene, we selected and used the probe with the most consistent pattern of regional variation across donors (i.e., differential stability) [112], calculated with:

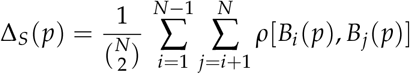

where *ρ* is Spearman’s rank correlation of the expression of a single probe, p, across regions in two donors *B*_*i* and *B*_*j*, and N is the total number of donors. Here, regions correspond to the structural designations provided in the ontology from the AHBA.

The MNI coordinates of tissue samples were updated to those generated via non-linear registration using the Advanced Normalization Tools (ANTs; https://github.com/chrisfilo/alleninf). Samples were assigned to brain regions by minimizing the Euclidean distance between the MNI coordinates of each sample and the nearest surface vertex. Samples where the Euclidean distance to the nearest vertex was more than 2 standard deviations above the mean distance for all samples belonging to that donor were excluded. To reduce the potential for misassignment, sample-to-region matching was constrained by hemisphere and gross structural divisions (i.e., cortex, subcortex/brainstem, and cerebellum, such that e.g., a sample in the left cortex could only be assigned to an atlas parcel in the left cortex) [49]. If a brain region was not assigned a sample from any donor based on the above procedure, the tissue sample closest to the centroid of that parcel was identified independently for each donor. The average of these samples was taken across donors, weighted by the distance between the parcel centroid and the sample, to obtain an estimate of the parcellated expression values for the missing region. All tissue samples not assigned to a brain region in the provided atlas were discarded.

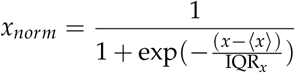

where ⟨*x*⟩ is the median and IQR_*x*_ is the normalized interquartile range of the expression of a single tissue sample across genes. Normalized expression values were then rescaled to the unit interval:

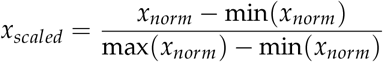

Gene expression values were then normalized across tissue samples using an identical procedure. Samples assigned to the same brain region were averaged separately for each donor and then across donors, yielding a regional expression matrix. We then excluded the right hemisphere regions due to the sparsity of samples and large number of regions with no expression data.

#### Transcriptomics maps of excitatory and inhibitory neuronal subtypes

The RNA sequencing of isolated cortical neuronal nuclei and their data-driven clustering has previously identified 16 neuronal subtypes, consisting of eight excitatory and eight inhibitory subtypes [44]. We used the list of subtype-specific genes and estimated the spatial distribution of each subtype by taking the weighted average mRNA expression of genes associated with the subtype, weighting each gene by the fraction of its positive values using exon-only derived transcripts per million [39]. Of note, few of the genes associated with each neuronal subtype had no eligible expression data in the processed AHBA dataset and were omitted from the calculation of subtype-specific gene expression (Table S1).

#### Correlated gene expression matrix

The correlated gene expression matrix was computed by correlating the regional gene expression patterns between the regions and across all genes, followed by its Z-transformation.

#### Partial least squares regression of LTC G1 with gene expression maps and developmental enrichment analysis

We used partial least squares regression (PLS) to identify genes that show high alignment in their regional expression variability with the LTC G1 within the left hemisphere. The scikit-learn package (https://scikit-learn.org/stable/) [114] was used to perform single-component PLS between the parcellated map of LTC G1 on one side, and the regional expression of all genes on the other side. We then ranked the genes based on the absolute value of their weights and selected the top 500 genes, which were subsequently divided into genes with positive and negative weight. Next, we fed the list of top genes with positive and negative alignment to LTC G1 to the online cell-type specific expression analysis (CSEA) developmental expression tool (http://genetics.wustl.edu/jdlab/csea-tool-2/) [53], which involves comparison of the genes against developmental expression profiles from the BrainSpan Atlas of the Developing Human Brain (http://www.brainspan.org), and for each developmental stage and brain structure reported the inverse log of Fisher’s Exact p-values corrected using Benjamini-Hochberg approach and based on the specificity index (pSI) threshold of 0.05.

### Dimensionality reduction of matrices

We applied the gradients approach implemented in the BrainSpace toolbox to identify the main axes (gradients) along which cortical regions can be ordered with regard to their similarity in the input matrix [33]. In this approach, to reduce the influence of noise on the embedding, the matrix is first sparsified according to the parameter p (default: 0.9), by zeroing out the p lowest-ranking cells in each row of the matrix. Next, the normalized angle similarity kernel function is used to compute the affinity matrix. Subsequently, principal component analysis, a linear dimensionality reduction technique is applied to the affinity matrix to estimate the macroscale gradients. Of note, to evaluate the robustness of our findings to analytical choices, we repeated this approach with alternative values of sparsity, as well as other, non-linear dimensionality reduction techniques including Laplacian eigenmaps and diffusion map embedding. As the signs of gradient values are arbitrary and for consistency in interpretation, the gradients in these alternative configurations were flipped if needed to match the sign of the original gradient values. The gradients approach was performed on the matrices of LTC, MPC, and structural covariance. In addition, the gradients approach was applied to the fused matrices of laminar thickness covariance and laminar density covariance. Following a previous work [76], the matrix fusion was performed by rank-normalizing both matrices, followed by rescaling of laminar density covariance ranks to that of laminar thickness covariance, and then horizontally concatenating the matrices.

### K-means clustering

We additionally used K-means clustering on the relative laminar thickness data to create a discrete map of laminar structure variability, as an alternative to the continuous map created using the gradients approach. The optimal number of clusters was identified using the yellowbrick package [115], by iteratively increasing the number of clusters, measuring the distortion score for each number of clusters, and identifying the elbow, after which adding more clusters does not considerably improve the model performance. The K-means clustering was performed using the scikit-learn package [114].

### Matrix associations

#### Matrix correlations

The Pearson correlation of any two given matrices was calculated after realigning the matrices to each other and removing the edges that were undefined in either matrix, across the edges in the lower triangle of the matrices. The null hypothesis testing for matrix correlations were performed non-parametrically, by creating a null distribution of correlation values calculated after random spinning of the parcels in one of the matrices for 1000 permutations. The spinning of parcels was performed using the ENIGMA Toolbox [46,116,117].

#### Association to geodesic distance

The effect of geodesic distance on continuous matrices was evaluated using an exponential fit, and its statistical significance was assessed based on pseudo R2 and using spin tests, similar to the approach used for matrix correlations.

#### Association of LTC to the cortical types

The average LTC for pairs of parcels within the same cortical type was calculated and compared to the average LTC for pairs of parcels that belonged to different cortical types using permutation testing. In each permutation (n = 1000) we spun the parcels in the LTC matrix, as described above, and calculated the difference of average LTC for the edges in the same versus different cortical types (for each cortical type, as well as across all cortical types), resulting in the null distribution of LTC differences within/between cortical types, which was used to calculate the p-values.

#### Association of LTC and geodesic distance to the connectivity probability

The structural connectivity matrix was binarized and logistic regressions were used to evaluate how connectivity probability relates to LTC and geodesic distance. The logistic regressions were performed using statsmodels package (https://www.statsmodels.org/stable/index.html) [118]. In each model pseudo R2 was reported and its statistical significance was assessed non-parametrically, using 1000 spin permutations of the LTC or geodesic distance matrix, as described above. The continuous changes in probability of connectivity as a function of LTC and geodesic distance was visualized by segmenting all the edges into 200 non-overlapping windows, sorted by the value of predictor, and plotting the probability of connectivity within each window, which was calculated by dividing the number of connected edges by the total number of edges within the window.

### Surface associations

#### Correlation of continuous maps

Brain regions that are closer tend to be more similar in their features compared to spatially distant regions, due to the spatial autocorrelation [49,50]. In null-hypothesis testing of surface data correlations it is important to take the spatial autocorrelation into account, and evaluate the correlation coefficients against a null model in which the spatial autocorrelation is preserved [50]. Therefore, we assessed the statistical significance for the correlation of surface maps using BrainSMASH (Brain Surrogate Maps with Autocorrelated Spatial Heterogeneity) (https://brainsmash.readthedocs.io/en/latest/) [50,119]. In this approach, surrogate surface maps are simulated with spatial autocorrelation that is matched to spatial autocorrelation in the original surface map, through creating random maps whose variograms are approximately matched to that of the original map. Of note, a few number of the reported correlations were performed between unparcellated data and at the level of vertices, and for these cases, we used an alternative approach of creating surrogates that preserve spatial autocorrelation, namely, by randomly spinning the sphere representation of the cortical mesh using the brainspace toolbox [33,116]. Subsequently, for the statistical testing of the correlation between surface maps X and Y, we generated 1000 surrogates of X, and created a null distribution by calculating the correlation coefficient of each X surrogate with the original Y, and compared the original correlation coefficient against this null distribution to calculate the p-value.

#### Association of continuous and categorical surface maps

The association of categorical surface maps with the continuous maps was assessed using one-way ANOVA, combined with post-hoc independent T tests which were corrected for multiple comparisons using Bonferroni correction. These tests were performed using spin permutation, by spinning the parcels of the continuous map and creating null distributions based on 1000 spun surrogates. The spinning of parcels was performed using the ENIGMA Toolbox [46,116,117]. To visualize continuous-categorical associations we either plotted the proportion of each category within each bin of the continuous variable, or used raincloud plots [120].

## Data and code availability

All the code and data for this study are openly available at a Github repository (https://github.com/amnsbr/laminar_organization). Our code, data and computing environment are published in a Docker image (https://hub.docker.com/r/amnsbr/laminar_organization), which can be used to reproduce our results and to perform additional analyses on the BigBrain data without having to install dependencies. The analyses in this project were performed predominantly using Python (version 3.9). BigBrain maps of cortical layers are available at https://ftp.bigbrainproject.org/. Other data used were either openly available online or acquired by contacting the authors and can be accessed in the project Github repository and Docker image. We refer the reader to the text and the project Github repository for the description and source of this data.

## Acknowledgements

The authors would like to thank the various contributors to the open-access databases that was used in our study. We would like to acknowledge the teams at the Foschungszentrum Julich and the Montreal Neurological Institute involved in making the BigBrain dataset available. Furthermore, we thank Dr. Budhachandra S. Khundrakpam and Prof. Alan C. Evans for providing the maturational coupling data. The authors would also like to acknowledge the financial support provided by various funders. This work was funded in part by Helmholtz Association’s Initiative and Networking Fund under the Helmholtz International Lab grant agreement InterLabs-0015, and the Canada First Research Excellence Fund (CFREF Competition 2, 2015–2016) awarded to the Healthy Brains, Healthy Lives initiative at McGill University, through the Helmholtz International BigBrain Analytics and Learning Laboratory (HIBALL), supporting A.S., C.P., S.B.E., B.C.B. and S.L.V.. S.L.V. and A.S. were additionally funded by the Max Planck Society (Otto Hahn award). K.W. was supported by the Wellcome Trust (215901/Z/19/Z). M.D.H. was funded by the German Federal Ministry of Education and Research (BMBF) and the Max Planck Society. B.C.B. acknowledges support from the SickKids Foundation (NI17-039), the National Sciences and Engineering Research Council of Canada (NSERC; Discovery-1304413), CIHR (FDN-154298), Azrieli Center for Autism Research (ACAR), an MNI-Cambridge collaboration grant, and the Canada Research Chairs program.

