## Supplementary Material for "The spatial arrangement of laminar thickness profiles in the human cortex scaffolds processing hierarchy"

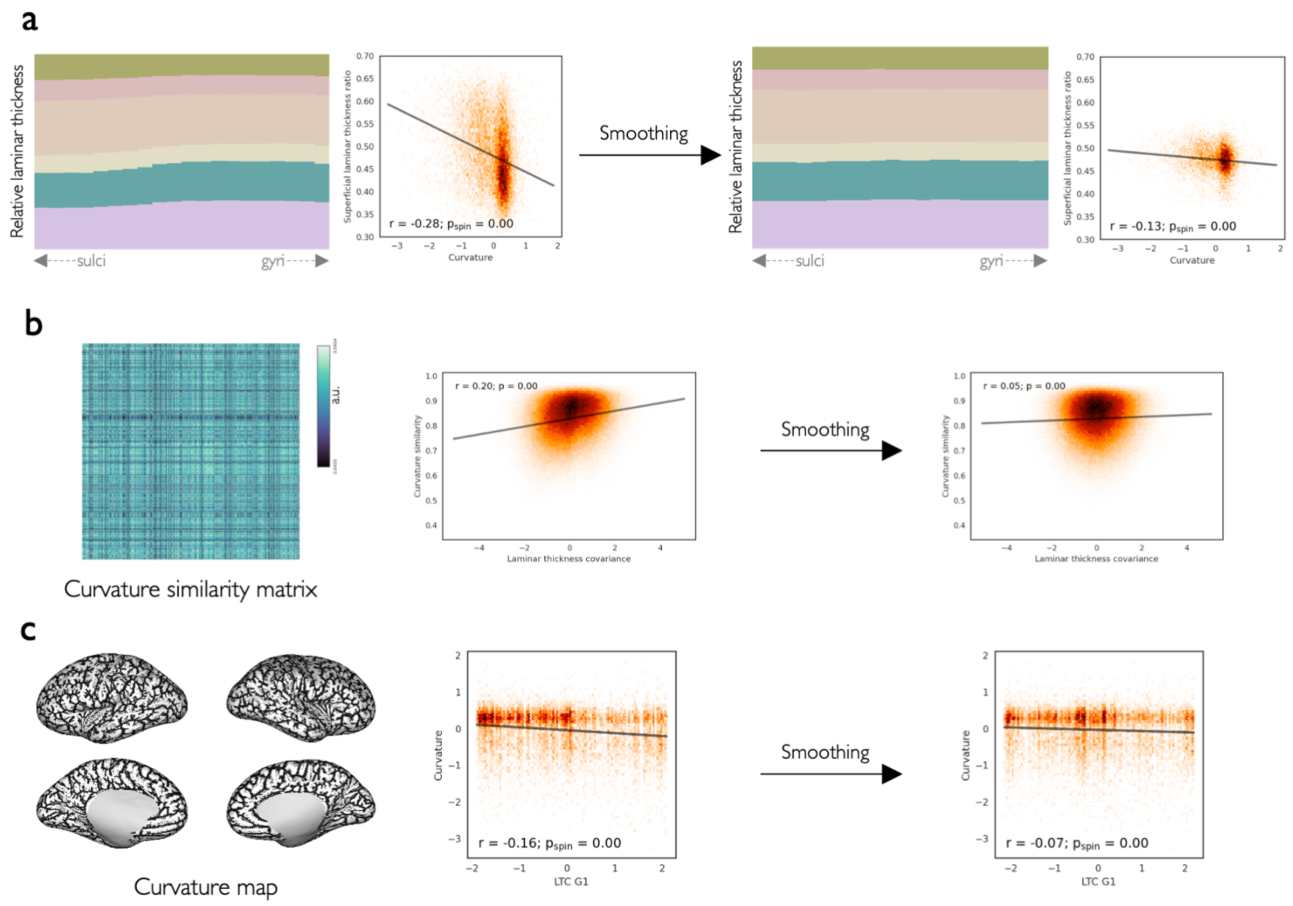


**Fig. S1. Effect of curvature on laminar thickness and laminar thickness covariance before and after smoothing. a)** The relative thickness of superficial layers decreases from sulci (negative curvature) to gyri (positive curvature) (*left*). After smoothing of the laminar thickness maps, the effect of curvature on laminar thickness was reduced remarkably, and the correlation of curvature with the relative thickness of superficial layers decreased (*right*). **b)** The matrix shows the similarity of parcels in their distribution of curvature values based on Jensen-Shannon divergence (*left*). The correlation of curvature similarity matrix with the LTC matrix decreased after smoothing (*right*). **c)** The curvature map (*left*) was significantly correlated with LTC G1, but the effect decreased after smoothing (*right*).

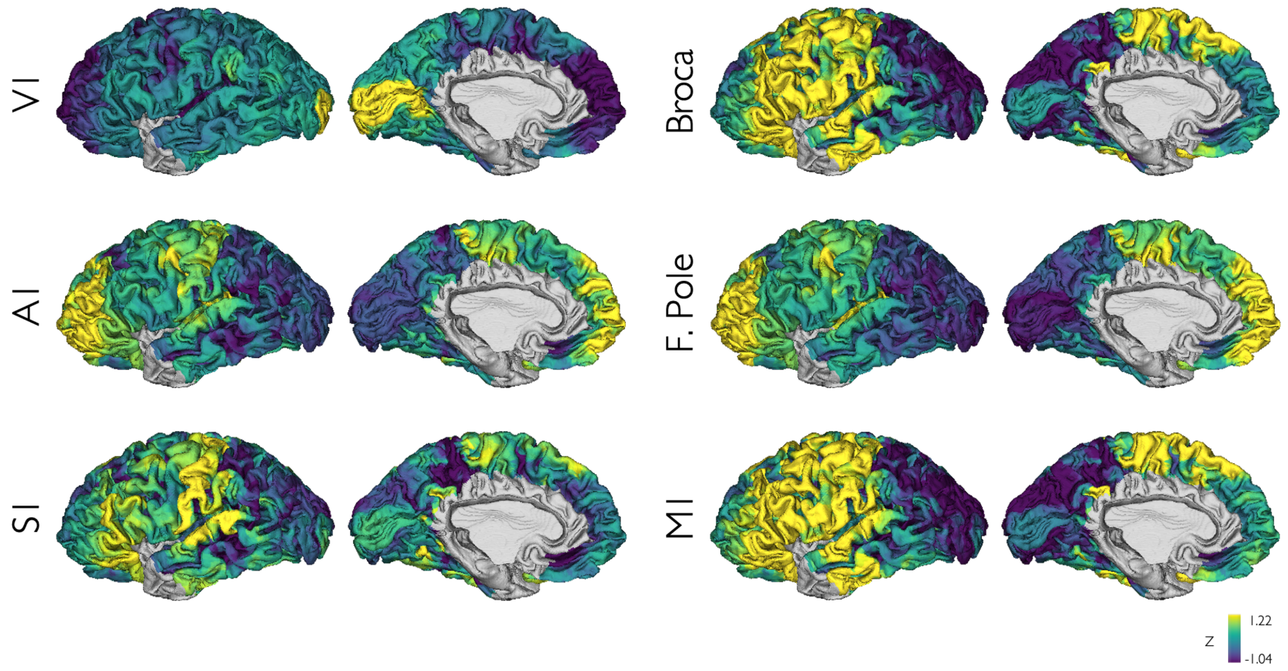


**Fig. S2. Laminar thickness covariance for regions of interest.** The laminar thickness covariance maps are shown for the centroid vertex of selected regions including the left primary visual cortex (V1), primary auditory cortex (A1), primary somatosensory cortex (S1), Broca's area, frontal pole and primary motor cortex (M1).

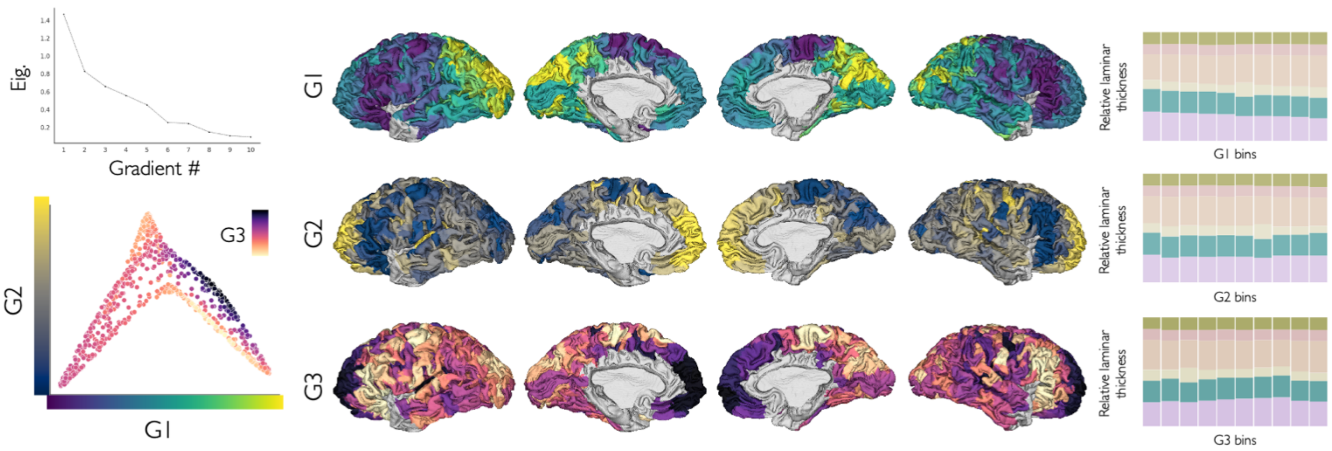


**Fig. S3. The first three axes of laminar thickness covariation.** *left top*: The first three gradients collectively explained 63.7% of the variance in LTC. *left bottom:* The scatter plot shows the position of brain regions in the gradient space of G1, G2 and G3. *center:* LTC G1, G2 and G3 projected on cortical surface show regional variation of laminar thickness across different axes. *right:* The pattern of relative laminar thickness variation along the three main axes.

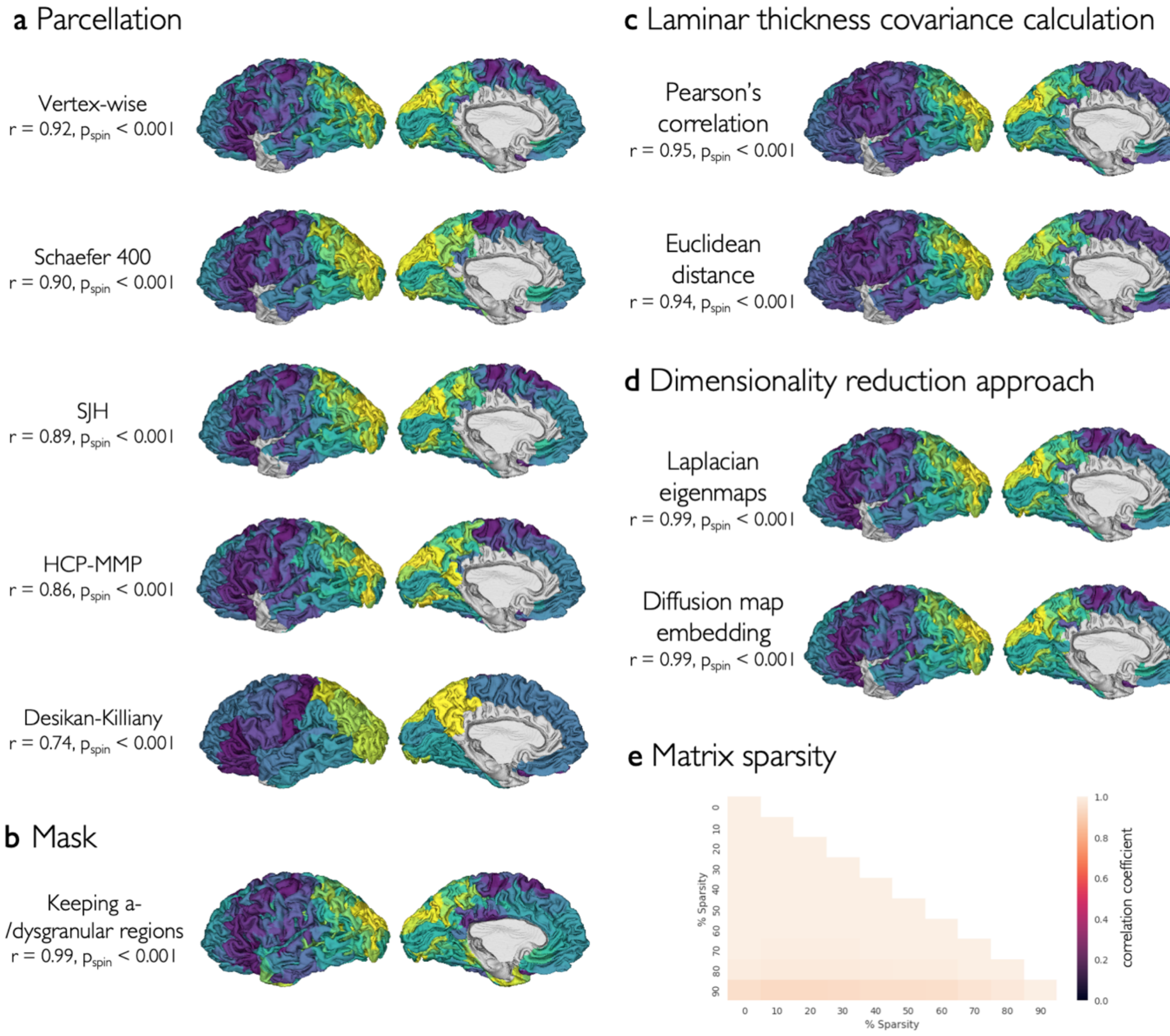


**Fig. S4. Robustness of LTC G1 to analytical choices.** LTC G1 spatial map was robust to the analytical choices. **a-d)** The maps of LTC G1 (left hemisphere) created using alternative analytical choices and their correlation with the original gradient are shown. **e)** The correlation of gradients created using different degrees of sparsity applied to the LTC matrix, from 0 to 0.9.

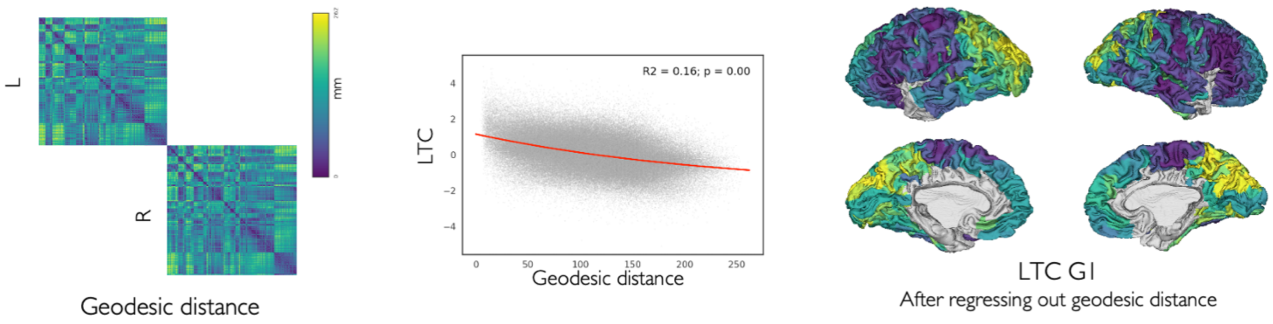


**Fig. S5. Association of laminar thickness covariance with geodesic distance.** Geodesic distance (*left*) showed an inverse exponential relationship with the LTC ($R^2 = 0.16$, $p_{spin} < 0.001$), indicating similar laminar thickness patterns between neighbor regions (*center*). The main axis of LTC G1 after regressing out the effects of geodesic distance (*right*) was significantly correlated with the original LTC G1 ($r = 0.97$, $p_{variogram} < 0.001$), indicating robustness of LTC G1 to geodesic distance.

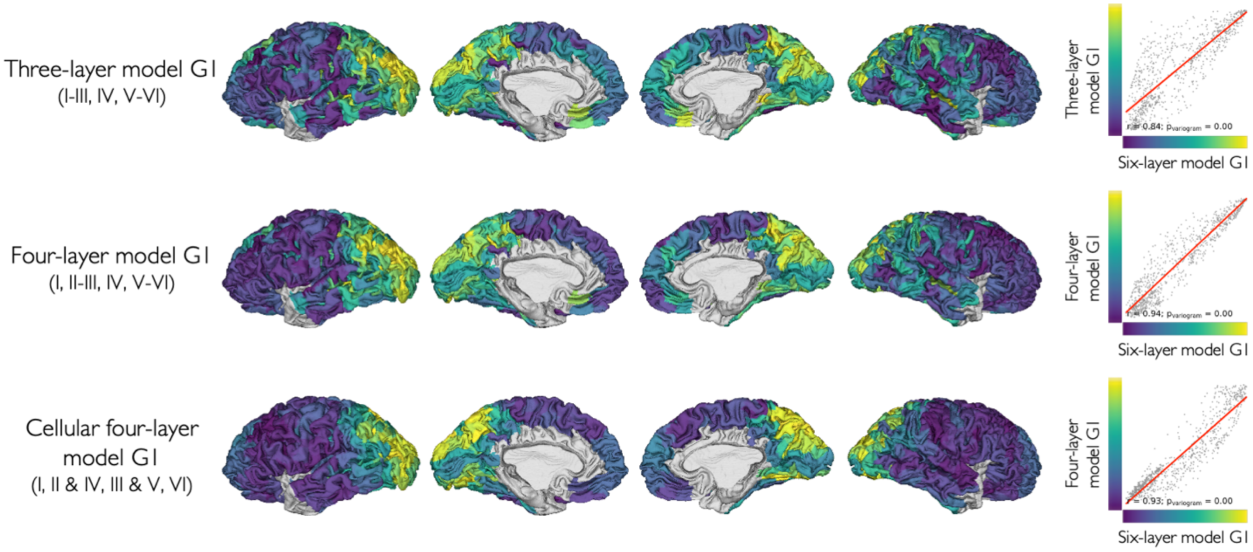


**Fig. S6. Models of laminar thickness variation assuming alternative laminar categorizations.** The three-layer, four-layer, and cellular four-layer models were created by first merging selected layers, and then calculating LTC and LTC G1 on the reduced laminar thickness data. Scatter plots show the correlation of these reduced models with the original six-layer LTC G1.

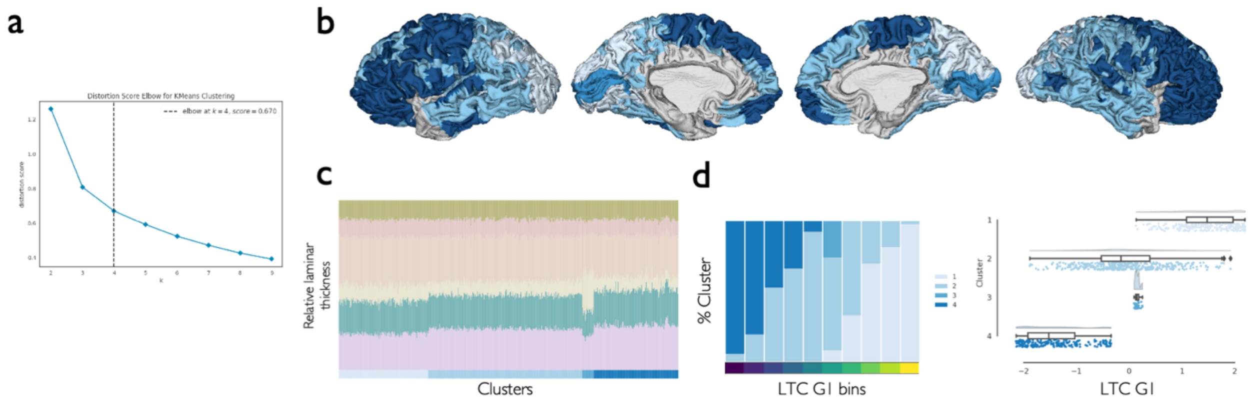


**Fig. S7. K-means clustering of cortical regions based on relative laminar thickness. a)** The distortion score of K-means clustering for the different number of clusters. The optimal number of clusters based on the elbow method was selected as four. **b)** Cluster of regions based on relative laminar thickness. **c)** Laminar thickness profiles of brain regions in each cluster. **d)** The LTC G1 values were significantly different between the clusters ($F = 813.1$, $p_{spin} < 0.001$). Post-hoc spin tests (Bonferroni-corrected) showed significantly different LTC G1 values between all pairs of clusters except 2 and 3.

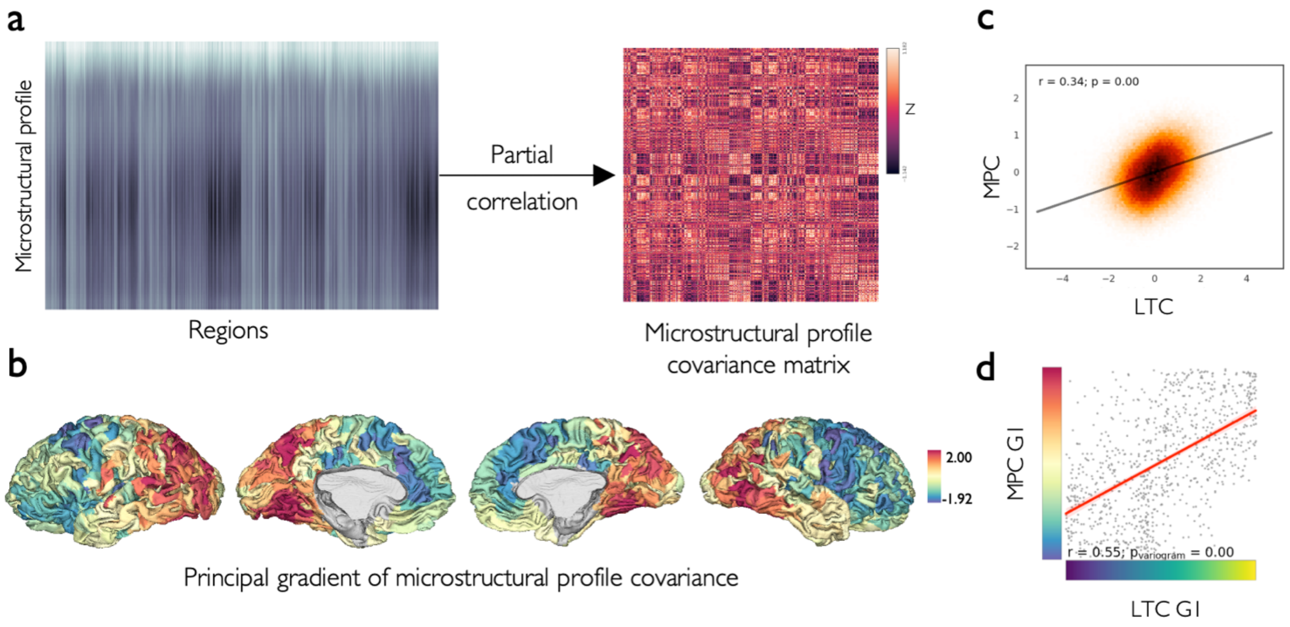


**Fig. S8. Microstructural profile covariance in association with laminar thickness covariance. a)** The average regional microstructural profiles show variations of BigBrain image intensity across cortical depth (50 samples). Microstructural profile covariance (MPC) matrix was created by the pairwise partial correlation of intensity profiles between the parcels. **b)** The principal axis of MPC created using principal component analysis. **c)** The correlation between MPC and LTC matrices. **d)** The correlation between main axes of LTC and MPC.

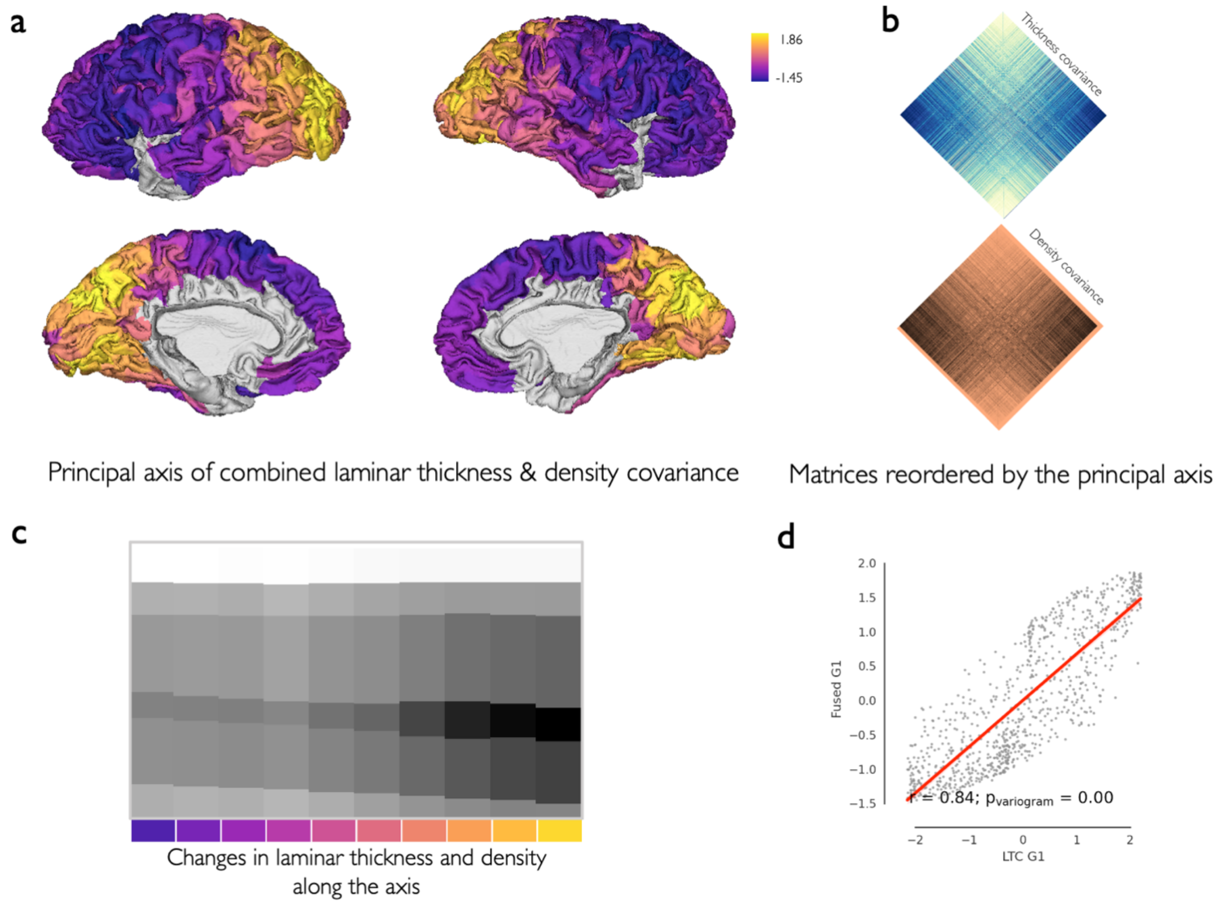


**Fig. S9. The principal axis of combined laminar thickness and density covariance. a)** The principal axis of combined laminar thickness covariance and laminar density covariance matrices (LTC-LDC G1). **b)** Laminar thickness and density matrices reordered by the LTC-LDC G1. **c)** The pattern of changes in the thickness and density of the six cortical layers along the LTC-LDC G1. **d)** The correlation of LTC-LDC G1 with MPC G1 and LTC G1.

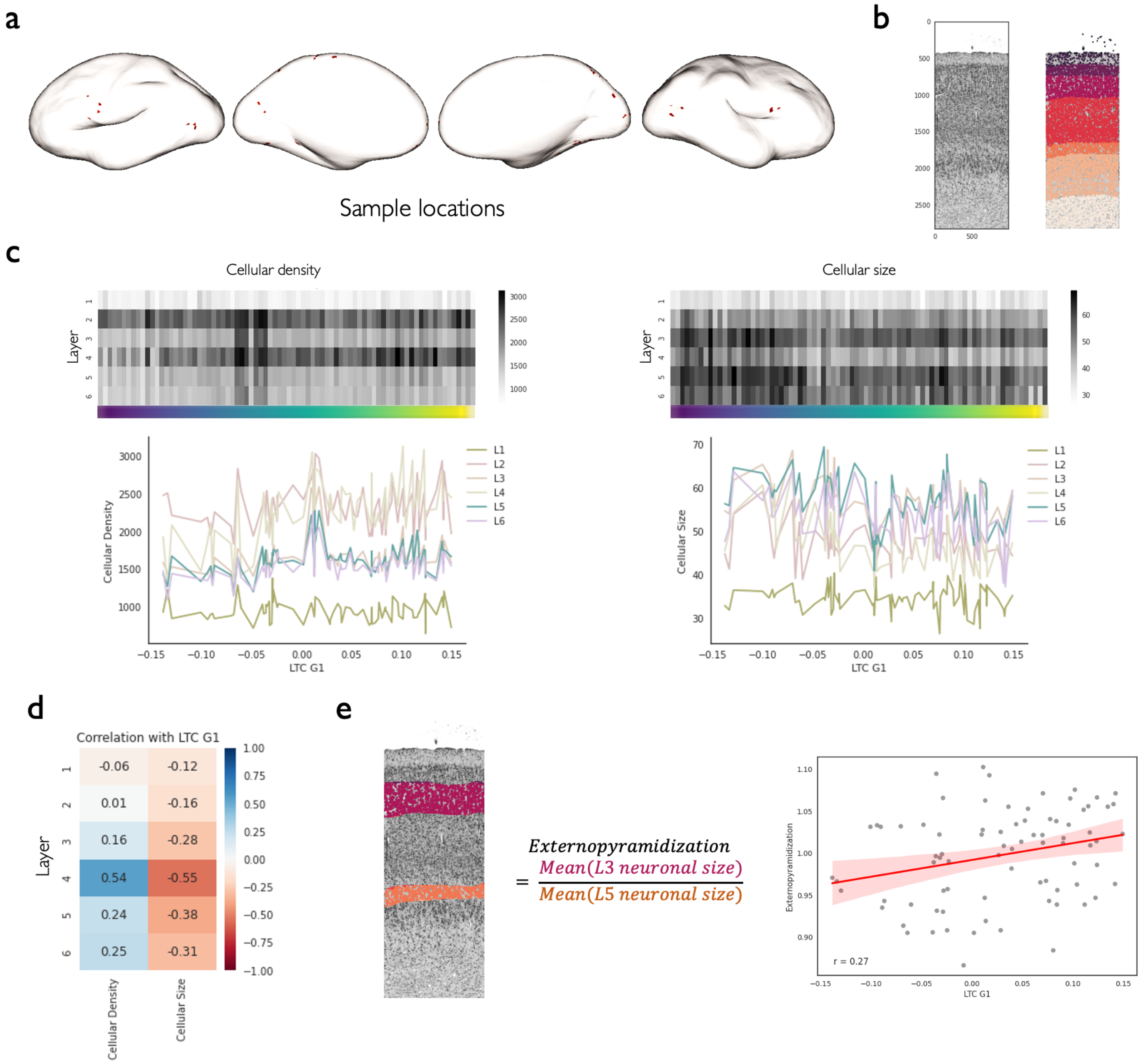


**Fig. S10. Laminar cellular features across the principal axis of laminar thickness covariance. a)** Locations of cortical samples for which laminar cellular data was available. **b)** Neuronal segmentation across cortical layers in an example sample. **c)** Variation of laminar neuronal density and size along the principal axis of laminar thickness covariation among the available samples. **d)** The correlation of laminar neuronal density and size along the LTC G1 among the available samples. **e)** The correlation of externopyramidization with the the LTC G1.

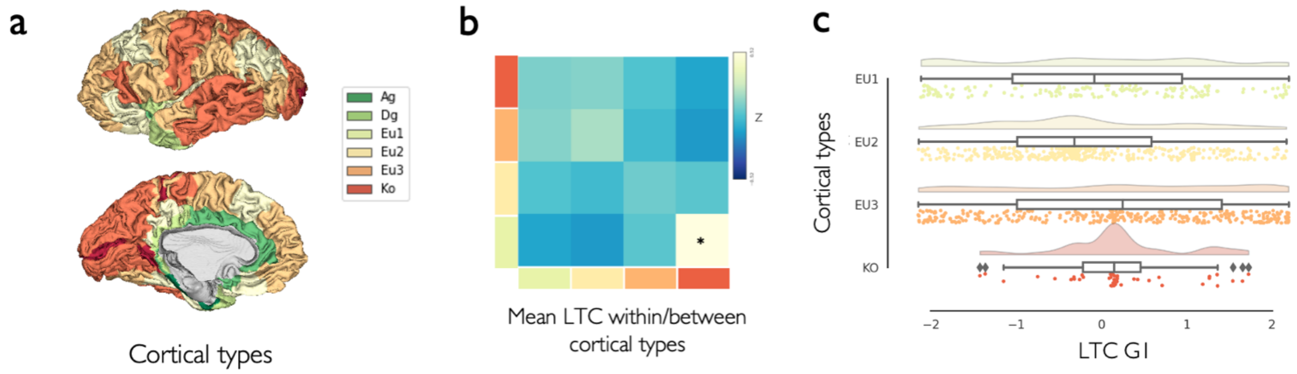


**Fig. S11. Association of cortical types with laminar thickness covariance. a)** The map of cortical types shows increasing laminar differentiation from agranular (green) to koniocortical (red) regions. **b)** The average LTC among pairs of parcels with the same or different cortical types, excluding agranular and dysgranular regions. Koniocortical regions showed significantly higher within-, compared to between-type average LTC. **c)** Distribution of LTC G1 across the cortical types are shown in a raincloud plot. No significant difference in LTC G1 values was observed between the cortical types ($F = 6.41$, $p_{spin} = 0.633$).

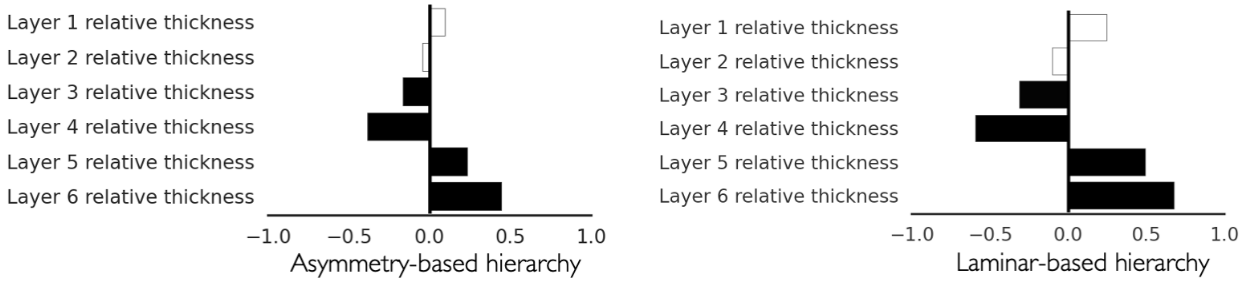


**Fig. S12. Association of asymmetry- and laminar-based hierarchy with the relative thickness of individual layers.** Bar length shows the correlation coefficient and its color represents the level of statistical significance from white ($p_{variogram, FDR} > 0.05$) to black ($p_{variogram, FDR} < 0.001$).

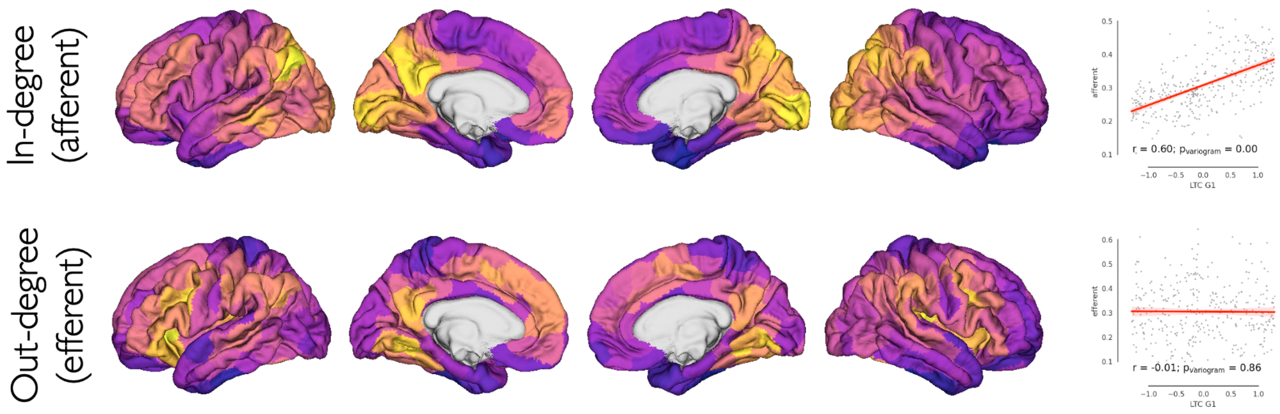


**Fig. S13. Afferent and efferent connectivity strength in association with LTC G1.** LTC G1 was significantly correlated with regional weighted in-degree (afferent strength) (*top*) but not weighted out-degree (efferent strength) (*bottom*).

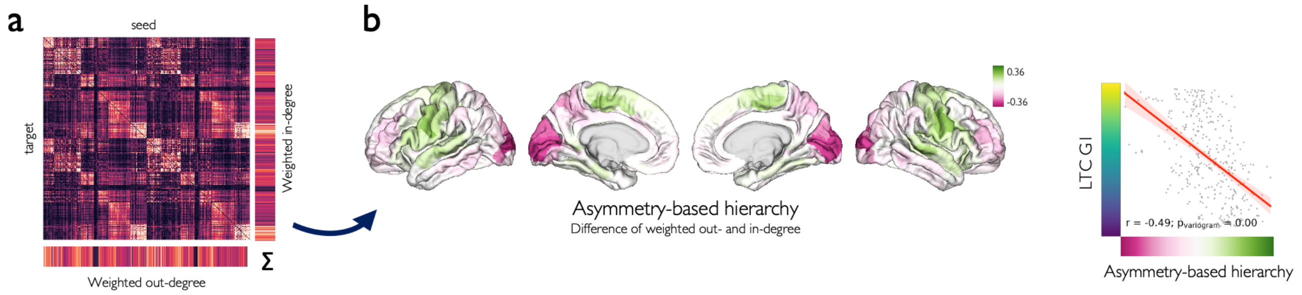


**Fig. S14. Association of LTC G1 with asymmetry-based hierarchy in the replication dataset. a)** The group-averaged effective connectivity matrix of the replication sample (N = 100) based on regression dynamic causal modeling. **b)** Regional asymmetry-based hierarchy was calculated as the difference between their weighted unsigned out-degree and in-degree, and was significantly correlated with LTC G1.

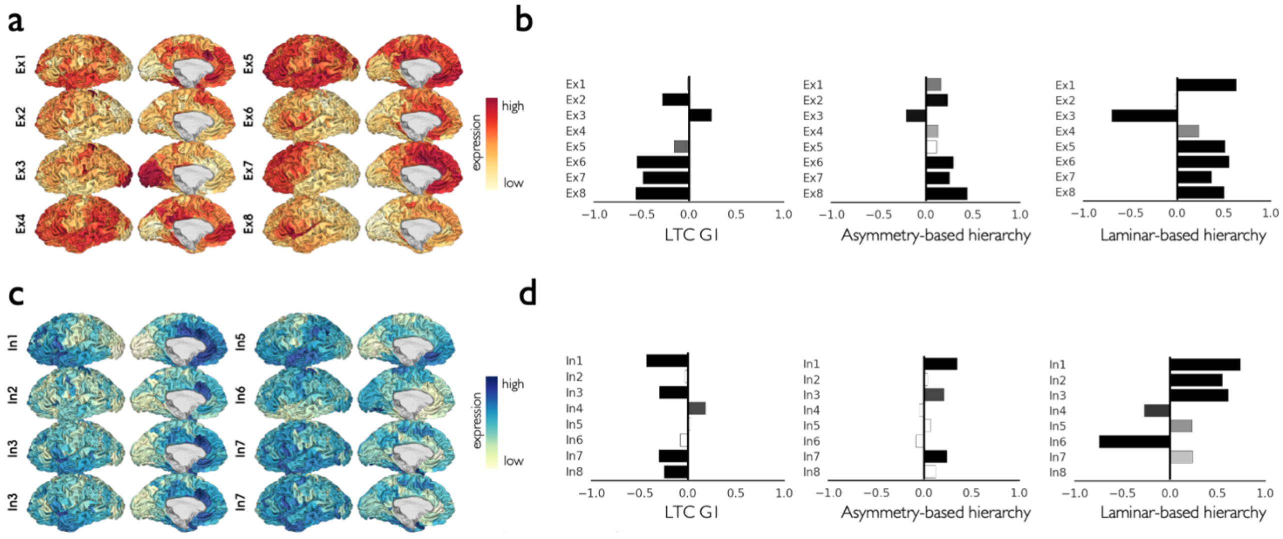


**Fig. S15. Microcircuitry in association with asymmetry-based hierarchy and LTC G1.** Spatial distribution of excitatory (a) and inhibitory (c) neuronal subtypes based on gene expression, and their correlation with LTC G1 as well as asymmetry-based and laminar-based hierarchy (b, d). Bar length shows the correlation coefficient and its color represents the level of statistical significance from white ($p_{variogram, FDR} > 0.05$) to black ($p_{variogram, FDR} < 0.001$).

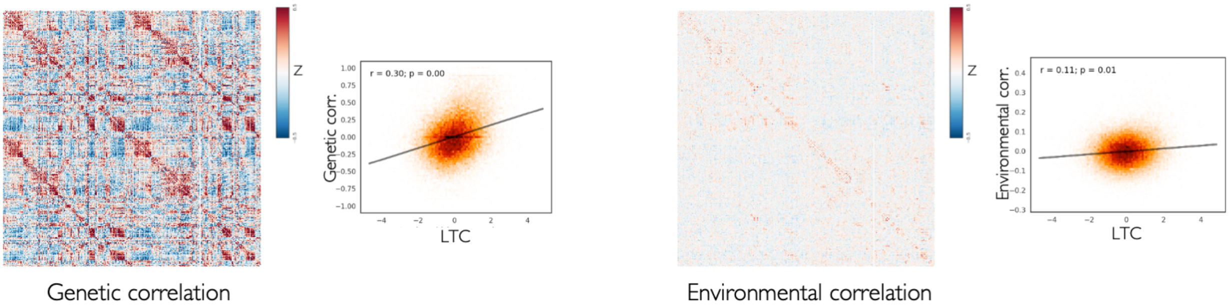


**Fig. S16. Inter-regional genetic and environmental correlation in association with laminar thickness covariance.** Inter-regional genetic and environmental correlation matrices based on cortical thickness in the HCP sample and their correlation with laminar thickness covariance.

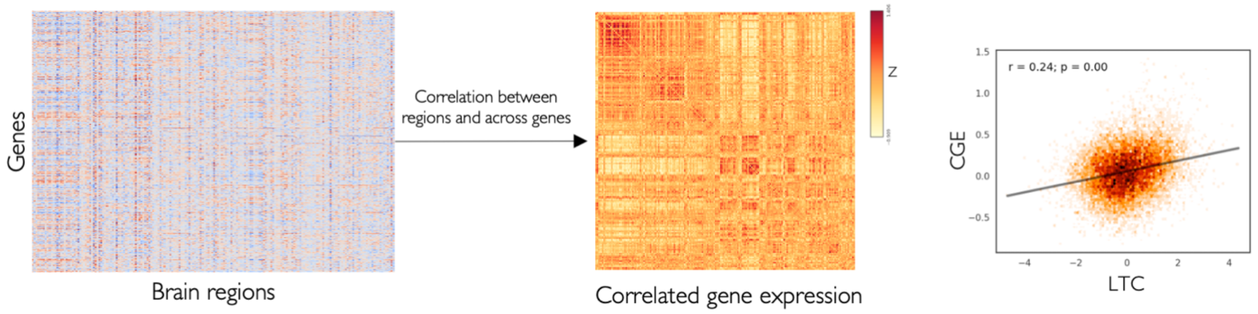


**Fig. S17. Correlated gene expression in association with laminar thickness covariance.** The gene expression data from Allen Human Brain Atlas (*left*) was used to create correlated gene expression (CGE) matrix, showing inter-regional similarity of gene expression profiles (*middle*), which was compared to the laminar thickness covariance matrix (*right*).

**Table S1. Category and number of genes associated with each neuronal subtype.**

| Neuronal subtype | Category | Number of genes<br>*Total (removed)* |
|---|---|---|
| **Excitatory** | | |
| Ex1 | *CPN* | 14 (0) |
| Ex2 | *GN* | 9 (1) |
| Ex3 | *GN* | 4 (0) |
| Ex4 | *SCPN* | 12 (1) |
| Ex5 | *SCPN* | 8 (0) |
| Ex6 | *SCPN* | 33 (0) |
| Ex7 | *CThPN* | 3 (0) |
| Ex8 | *CThPN* | 57 (5) |
| **Inhibitory** | | |
| In1 | *VIP+ RELN+ NDNF+* | 2 (0) |
| In2 | *VIP+ RELN- NDNF-* | 4 (0) |
| In3 | *VIP+ RELN+ NDNF-* | 12 (0) |
| In4 | *VIP- RELN+ NDNF+* | 6 (0) |
| In5 | *CCK+ nNOS+ CB+* | 7 (2) |
| In6 | *PV+ CRHBP* | 4 (0) |
| In7 | *SOM+ CB+ NPY+* | 6 (0) |
| In8 | *SOM+ nNOS+* | 1 (0) |

CPN: cortical projection neuron, GN: granule neuron, SCPN: subcortical projection neuron, CThPN: corticothalamic projection neuron, SOM: somatostatin, NPY: neuropeptide Y, CB: calbindin-D-28k, VIP: vasoactive intestinal peptide, RELN: reelin, nNOS: neuronal nitric oxide synthase, PV: parvalbumin, CCK: cholecystokinin, NDNF: neuron-derived neurotrophic factor; CRHBP: corticotropin releasing hormone binding protein.
